# An instructive role for IL7RA in the development of human B-cell precursor leukemia

**DOI:** 10.1101/2020.01.27.919951

**Authors:** Ifat Geron, Angela Maria Savino, Noa Tal, John Brown, Virginia Turati, Chela James, Jolanda Sarno, Yu Nee Lee, Avigail Rein Gil, Hila Fishman, Yehudit Birger, Inna Muler, Michal Hameiri-Grossman, Kara Lynn Davis, Victoria Marcu-Malina, Oren Parnas, Ute Fischer, Markus Müschen, Arndt Borkhardt, Ilan Richard Kirsch, Arnon Nagler, Tariq Enver, Shai Izraeli

## Abstract

B-cell precursor acute lymphoblastic leukemia (BCP-ALL) is preceded by a clinically silent pre-leukemia. Experimental models that authentically re-capitulate disease initiation and progression in human cells are lacking. We previously described activating mutations in interleukin 7 receptor alpha (IL7RA) that are associated with the poor-prognosis Philadelphia-like (Ph-like) subtype of BCP-ALL. Whether IL7RA signaling has a role in initiation of human BCP-ALL is unknown.

IL7RA is essential for mouse B-cell development; however, patients with truncating *IL7RA* germline mutations develop normal mature B-cell populations. Herein, we explore the consequences of aberrant IL7RA signaling activation in human hematopoietic progenitors on malignant B-cell development.

Transplantation of human cord-blood hematopoietic progenitors transduced with activated mutant IL7RA into NOD/LtSz-*scid IL2Rγ^null^* mice resulted in B-cell differentiation arrest with aberrant expression of CD34^+^ and persistence of pro-B cells that survive despite failing to achieve productive rearrangement of immunoglobulin V(D)J gene segments. Activation of IL7RA signaling enhanced self-renewal and facilitated the development of a BCP-ALL in secondary transplanted mice. The development of leukemia was associated with spontaneous acquired deletions in CDKN2A/B and IKZF1 similar to what is observed in Ph-like BCP-ALL in humans. Single cell gene expression analysis suggested that pre-leukemic cells resided within a subpopulation of early B-cell precursors with CD34^+^CD10^high^CD19^low^ immunophenotype.

The development of a bona fide BCP-ALL from IL7RA transduced cells supports the hypothesis that aberrant activation of the IL7RA pathway in human B-cell lineage progenitors has an instructive role by creating a pre-leukemic state which is vulnerable to transformation. These are the first demonstrations of a role for IL7RA in human B-cell differentiation and of a de-novo Ph-like BCP-ALL development from normal human hematopoietic progenitors *in vivo*.

## Introduction

The current paradigm of the evolution of B-cell precursor acute lymphoblastic leukemia (BCP-ALL) suggests two distinct stages: A commonly occurring initiating genetic event that generates pre-leukemia and a rare progression to leukemia through the acquisition of additional somatic genetic events^1^. In childhood ALL the initiating event occurs most commonly in-utero and consists usually of an aberration of a transcriptional regulator. Progression to leukemia is caused by a series of acquired genetic aberrations that halt B-cell differentiation and increase cell proliferation, survival and self-renewal^2, 3^. Increased signaling through RAS or STAT5 pathways are typical progression events and are generally thought to act as the “fuel” enhancing leukemic cell growth^4–7^. However, activation of signaling may also initiate BCP-ALL, as in the case of *BCR-ABL1* translocation, that is generally perceived as a leukemia-initiating event. Yet, this assumption has never been proven experimentally in human hematopoietic progenitor cells.

Interleukin-7 receptor alpha (IL7RA) is a receptor subunit with dual roles. Upon association with Interleukin-2 receptor gamma (IL2Rγ) subunit, it forms the Interleukin-7 (IL7) receptor and when bound to cytokine receptor-like factor 2 (CRLF2) subunit it constitutes the thymic stromal lymphopoietin (TSLP) receptor. Loss-of-function mutations in IL7RA are associated with absent B and T-cells in mice but with the absence of only T-cells in humans^8, 9^. Thus, while IL7RA is important for mouse T and B lymphopoiesis its role in human B-cell development is unclear^10–12^.

Activation of IL7RA pathway is commonly associated with T-cell malignancies^13^. We previously described IL7RA activating mutations in Philadelphia-like (Ph-like) BCP-ALL^14^. “Ph-like” leukemia is a subgroup of BCP-ALLs that are associated with poor prognosis and characterized by a similar gene expression signature to *BCR-ABL1* ALL^15–17^. The majority of these leukemias are characterized by aberrant activation of the TSLP receptor and the downstream JAK/STAT signaling pathway^18^. Genomic studies have provided conflicting evidence regarding the role of CRLF2/IL7RA aberrations in leukemic evolution. Aberrant expression of the receptors may be clonal or subclonal, preserved or altered between diagnosis and relapse^7, 19–21^. Two mouse models suggest that under specific conditions, expression of CRLF2 and/or IL7RA may initiate BCP-ALL ^22–24^. Nevertheless, the relevance of these models to human BCP-ALL is unclear, due to major differences in the role of both IL7 and TSLP signaling in B-cell development in humans. Hence, the role of CRLF2/IL7RA in human leukemia initiation, if any, is unknown.

Here we provide the first experimental evidence in human hematopoietic cells that expression of activated IL7RA (IL7RAins), with or without CRLF2 has an instructive role in human B-cell development by initiating a pre-leukemic state that is vulnerable to evolve to overt “Ph-like” BCP-ALL.

## Methods

For detailed methodology see supplementary material

### Human CD34^+^ hematopoietic progenitors

Fresh cord blood (CB) units were obtained from Sheba Medical Center CB bank under Institutional Review Board–approved protocols to obtain CB units for research purposes (Approval 5638-08-SMC, anonymized units that are otherwise discarded due insufficient volume for public storage). CD34^+^ cells were isolated using magnetic beads (Miltenyi USA) following conventional method.

### IL7RA and CRLF2 overexpression

Cloning of IL7RA and CRLF2 into B-cell specific lentiviral vectors (Kindly provided by Rawlings lab^25^) was performed with standard cloning protocols (see supplementary material). Production of lentiviral vectors was done as previously described^26^, and used to transduce CD34^+^ hematopoietic progenitors.

### Xenotransplants

NOD/LtSz-scid IL2Rγnull (NSG) mice were purchased from Jackson laboratories (Mount Desert Island, Maine, USA). Mice were bred and housed in specific pathogen-free conditions. All animal experiments were approved by the Animal Care Committee at Sheba Medical Center (IRB 1007/15). 5-8-week-old NSG females were irradiated (1-1.5 Gy X-ray) 4-24 hours prior to transplantation. For primary transplantations, 1×10^5^-1.5×10^5^ cells were transduced 72-96 hours prior to transplantation. On transplantation days cells were sampled to assess transduction efficiency and transplanted via tail vain injection. 23-30 weeks post transplantation, mice were euthanized and hematopoietic tissues (spleen, bone marrow from femurs, liver and peripheral blood) were harvested.

### Flow cytometric analysis

Cells were stained with fluorochrome-conjugated antibodies using standard staining and analysis protocols. Data analysis was done using Kaluza software (Beckman-Coulter, California, USA)

### RNA/DNA sequencing and expression profile analysis

Bulk RNA sequencing: RNA was purified from 5000-20000 transduced CD45^+^CD3^−^ cells that were sorted from spleens of transplanted mice (see details in supplementary methods). cDNA libraries were prepared using SMARTer kit (Clontech Laboratories, Inc. A Takara Bio Company, Mountain View, CA, USA) followed by the Nextera protocols (Illumina, CA, USA). Genome-wide expression profiles were obtained by sequencing of the samples on Illumina NextSeq 500 using NextSeq 500/550 High Output v2 kit.

Single cell RNA sequencing (scRNAseq): 10X library was prepared from 4000-10000 cells (10X V3 library preparation kit, 10X, USA) that were sorted from BM of leukemic/pre-leukemic/BB engrafted mouse and sequenced on NextSeq 500 System (Illumina USA)

### B-cell Immune Repertoire sequencing

5000-20000 transduced CD45^+^ CD10^+^ and CD19^+^ engrafted cells were sorted. gDNA was purified using QIAamp® DNA Micro kit (Qiagen Inc. USA). B-cell repertoire sequencing was performed using Adaptive ImmunoSEQ IGH deep assay at Adaptive Biotechnologies (Seattle, WA, USA). Analysis of B-cell receptor repertoire was done by Adaptive Biotechnologies using proprietary pipeline^27^.

### Whole genome sequencing

Leukemic and BB transduced corresponding CB cells were collected from transplanted mice. Sequencing libraries were prepared using NEBNext ULTRA II library preparation kit (see details in supplementary methods) and sequenced on HiseqXten (BGI Hong Kong).

### Statistical analysis

Data was analyzed using Microsoft Excel and GraphPad Prism software (La Jolla, CA). Data is either depicted as mean ± SE or as a scatter plot with mean ± SE. Comparisons between groups were performed by unpaired student t-tests in two groups analysis, by one-way ANOVA tests when more than two groups were compared and groups had equal variance or in Kruskal-Wallis-test – a one-way non-parametric analysis of variance, when no equal variance could be assumed. Post hoc analyses were done either by using Dunnett post hoc analysis – to compare samples to control group or by Dunn’s/Tukeys multiple comparison test to compare between all experimental groups. P values < 0.05 were considered statistically significant.

## Results

### Activation of IL7RA pathway blocks differentiation of human B-cells at the progenitor stage *in-vivo*

To test the role of activated IL7RA in leukemia initiation we expressed wild type and/or an activated mutant form of human IL7RA containing an in-frame insertion PPCL (Ins12 (CCCCCGTGCCTA) position 243) (IL7RAwt/ins) in human CB hematopoietic progenitors. As IL7RA mutations in BCP-ALL frequently correlate with aberrant CRLF2 expression, combinations of IL7RA and CRLF2 were used. The coding sequences were cloned into a lentiviral vector with a bi-cistronic cassette under the expression control of an Eμ-B29 promoter/enhancer to augment expression in B-cell precursors. Backbone vector expressing GFP (BB) was used as control^25^ (supplementary figure 1A). Activity of the IL7RA/CRLF2 transgenes was verified by STAT5 phosphorylation assay in BCP-ALL cell line (Supplementary figure 1B). Transduced CB hematopoietic progenitors (CD34^+^) were transplanted into NOD/LtSz-*scid IL2Rγ^null^* (NSG) mice.

Development of BCP-ALL leukemia is associated with a block in B-cell differentiation at pro/pre-B-cell stage. We therefore analyzed the differentiation pattern of the human B-lineage 24-30 weeks post-transplantation. B-cell differentiation beyond the pre-B-cell stage (CD19^+^CD10^+^sIgM^−^) was significantly inhibited in activated IL7RA transduced cells (*Figure 1A-D*). To further define B-lineage differentiation stage of transduced cells, engrafted cells were analyzed by mass cytometry, and a single cell developmental classifier was applied as previously described by Good Z. et al.^28^. As depicted in supplementary figure 2A, an enlarged pre-BI population is observed in the CRLF2-IL7RAins transduced cells and an earlier pro-BII fraction in the IL7RAins alone transduced cells.

**Figure 1.**
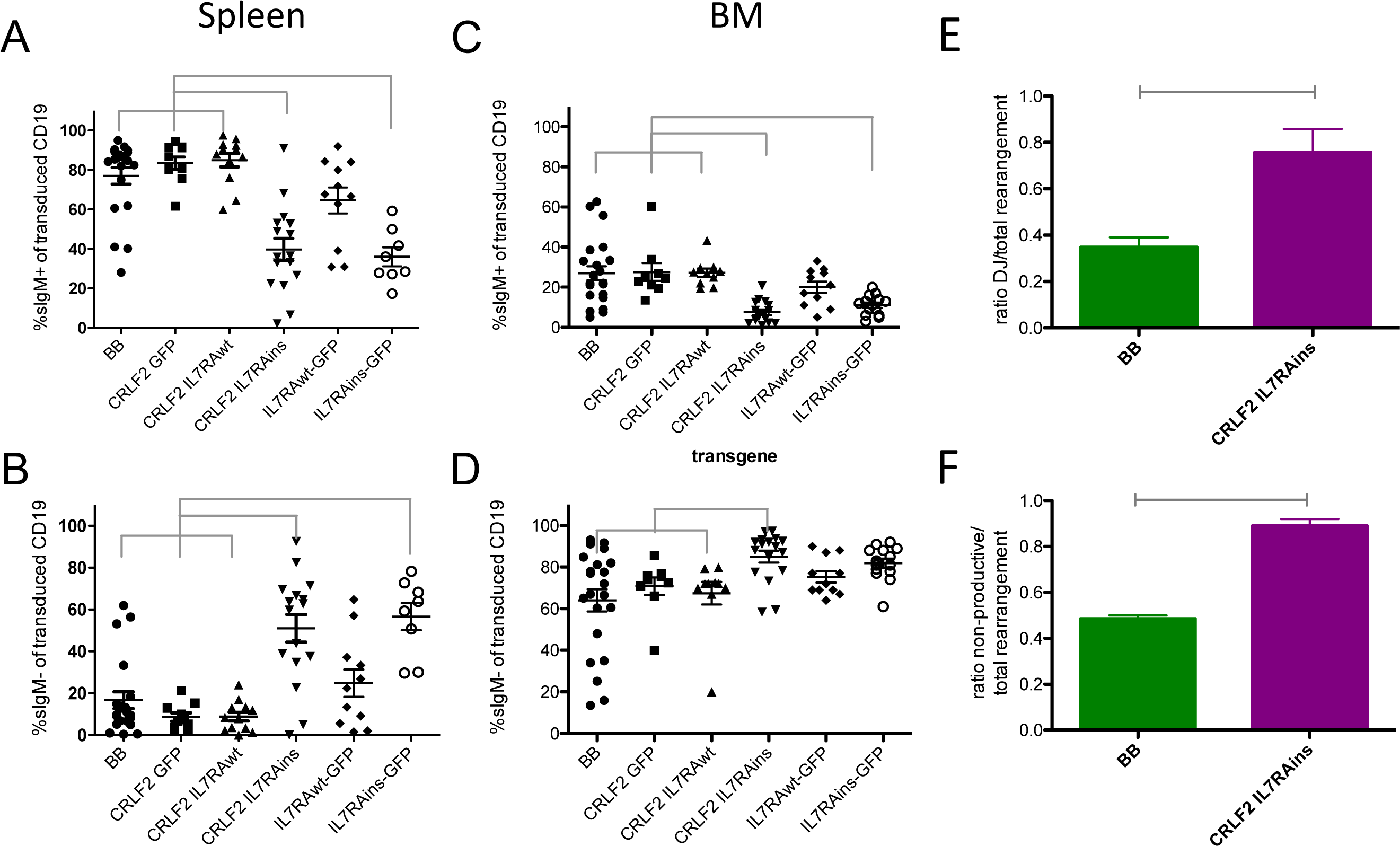
CRLF2/IL7RA transduction alters B-lineage differentiation of human CB CD34^+^ progenitors transplanted in immune-deficient mice. A-D B-lineage differentiation of human CB CD34^+^ cells expressing GFP (BB), CRLF2-GFP, CRLF2-IL7RAwt, CRLF2-IL7RAins, IL7RAwt-GFP and IL7RAins-GFP in spleen (A, B) and BM (C,D) of engrafted mice. A+C: differentiation to immature/naïve-B-cells (sIgM+); B+D: differentiation to pro-pre-B-cells (sIgM^−^). Dot plots show sample scatter with mean +/− SEM. Gray linkers indicate statistically significant difference (p<0.05) between groups. Statistical analyses were performed using Kruskal-Wallis non-parametric test with Dunn’s post-hoc analysis. E-F: V(D)J rearrangement analysis of CD10^+^ and CD19^+^ BB/CRLF2-IL7RAins transduced cells sorted from BM of transplanted mice. (E) Bar-graph representing fraction of DJ rearranged of the total rearranged IgH loci in transduced cells. (F) Bar-graph representing ratio of non-productive to productive rearrangement in transduced cells. Bars are mean +/− SEM of BB (n=3) and CRLF2-IL7RAins (n=4). Gray linkers indicate statistically significant difference (p<0.05) between groups. Statistical analyses were performed using two tailed t-test.

**Figure 2.**
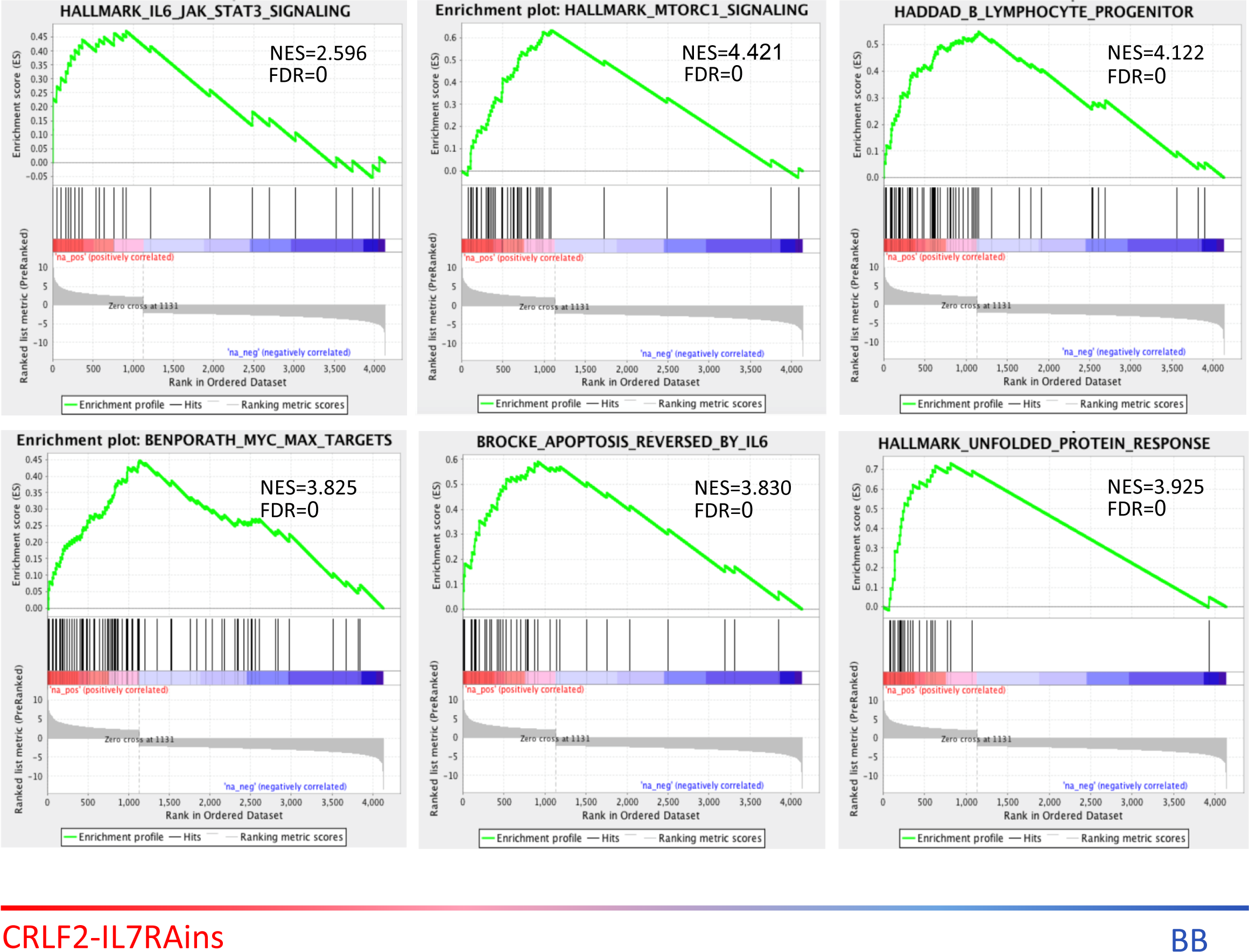
Signaling activation in engrafted CRLF2-IL7RAins transduced cells. GSEA plots of differentially expressed genes between transplanted CRLF2-IL7RAins and BackBone transduced cells.

Since the stage of B-cell differentiation is reflected in a cell’s V(D)J recombination status, we performed B-cell repertoire sequencing of the IgH locus in sorted transduced CD10^+^CD19^+^ cells from bone marrow (BM) of transplanted mice. Consistent with the early B-cell differentiation stage that was observed by immunophenotyping, the fraction of DJ rearranged cells was significantly expanded in the CRLF2-IL7RAins transduced population (figure 1E). Clonality analysis (supplementary figure 2B) did not show significant expansion of specific clones suggesting that the accumulation of B-cell precursors was due to a general block in differentiation and not to enhanced growth of specific clones.

During normal B-cell differentiation, cells carrying non-productive V(D)J rearrangements undergo programmed cell death^29, 30^. In contrast, acute lymphoblastic leukemia cells often carry non-productive V(D)J rearrangements^31^. We observed a substantial increase in the ratio of the non-productive rearranged fraction in the CRLF2-IL7RAins transduced cells (figure1F). This observation suggests that activation of TSLP/IL7RA signaling provided an enhanced survival capacity of the cells that otherwise would have been destined to programmed death in the absence of productive B-cell receptor rearrangements.

### Gene expression analysis of CRLF2/IL7RA activated xenografted cells reveals activation of JAK-STAT, mTOR and survival pathways

To decipher the mechanism underlying the observed B-cell phenotypic changes we compared the gene expression profile of CRLF2-IL7RAins transduced human cells to that of BB transduced controls from the spleens of primary engrafted mice. As expected, CRLF2 and IL7RA were ranked high in the list of differentially expressed genes confirming successful transduction (supplementary table 1). As can be anticipated after activation of the CRLF2/IL7RA signaling^32^, enrichment of gene sets representing JAK-STAT and mTOR signaling were observed (figure 2). Consistent with the phenotypic B-cell differentiation block the signatures were enriched with B-cell precursor gene expression and with higher expression of RAG1 and RAG2 compared to the BB transduced group (supplementary table 1).

In agreement with their aberrant survival at the presence of non-productive immunoglobulin heavy chain gene V(D)J gene rearrangements, transduced cells also displayed significant enrichment for gene sets associated with survival/proliferation pathways (MYC pathway, rescue of apoptosis by IL6 signaling – figure 2). Furthermore, enrichment of the unfolded protein response gene set, with upregulation of XBP1 (FC 1.95) and HSPA5 (FC 1.42) that were previously demonstrated to be essential for pre-B and pre-B ALL cells survival^33^ were detected (figure 2 and supplementary table 2). Thus, gene expression analysis demonstrated activation of pathways promoting the survival of transduced B-cell progenitors.

### Aberrant expression of activated IL7RA induces a pre-leukemia B-cell precursor immunophenotype retaining self-renewal capacity

B-cell precursor leukemic cells express the hematopoietic progenitor marker CD34 that is normally silenced past the early-B-cell progenitor differentiation stage^34, 35^. Consistent with the enrichment in early B-cell progenitors (supplementary figure 2A), activation of IL7RA pathway resulted in expansion of the CD19^+^CD10^+^CD34^+^ population (figure 3A). In six out of 47 mice engrafted with activated-IL7RA transduced cells, we have identified a unique CD10^high^CD19^low^ sub-population that was undetectable in control groups (figure 3B and supplementary figure 3). This population was enriched with CD34^+^ cells and can either represent an expanded early-B population or a distinct “pre-leukemic” population.

**Figure 3.**
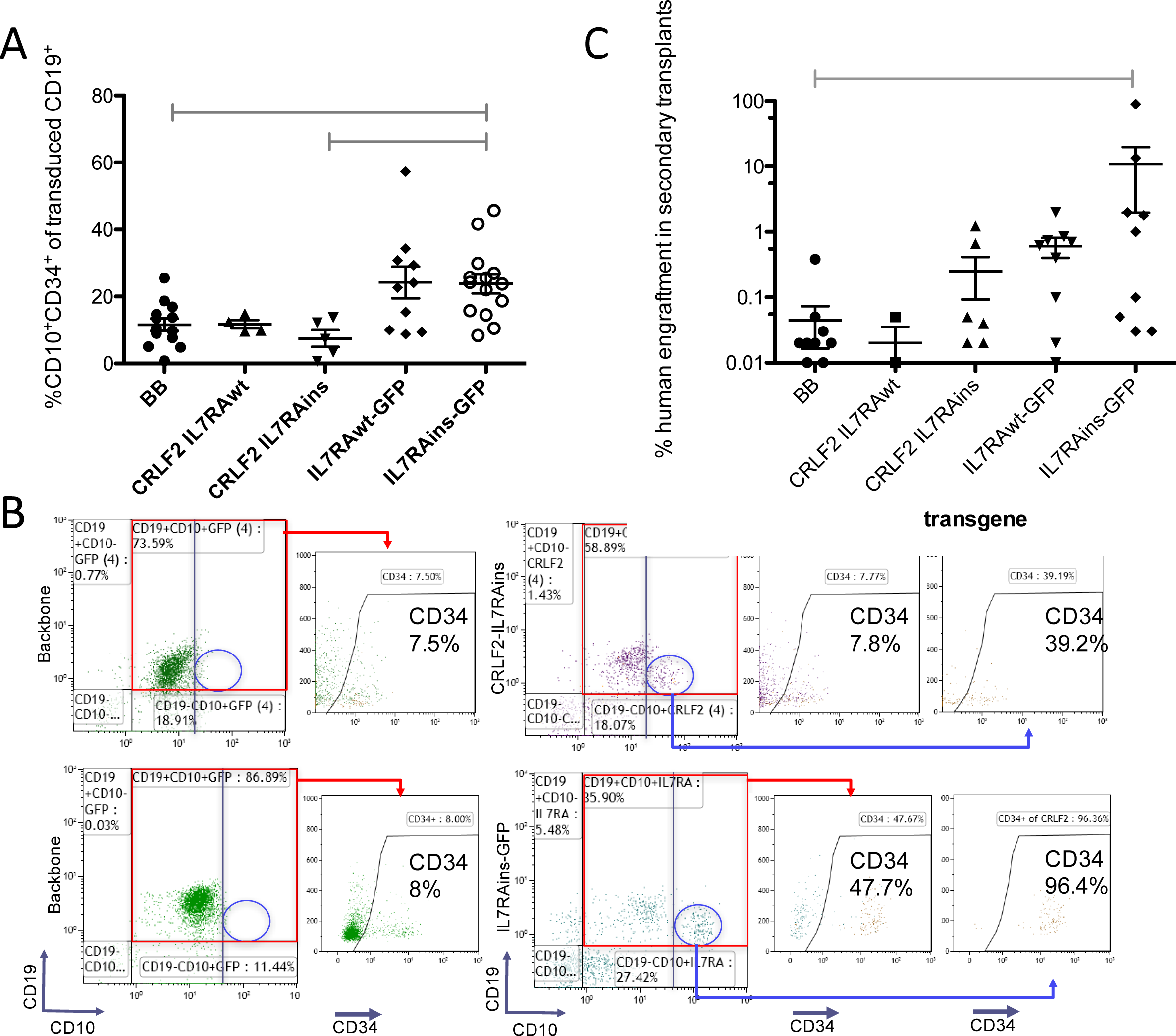
Enhanced CD34^+^CD10^+^ expression and self-renewal of IL7RA activated cells. (A) Relative CD10^+^CD34^+^ population of engrafted transduced human CD19 cells in BM (BB n=11, CRLF2-IL7RAwt n=4, CRLF2-IL7RAins n=5, IL7RAwt-GFP n=10, IL7RAins-GFP n=14). (B) Flow cytometry immunophenotyping of engrafted backbone and CRLF2-IL7RAins or IL7RAins transduced cells. Samples in the same row are from the same CB batch. Arrows indicate that the gated population was analyzed in the following scatters. (C) Percentage of human cells in BM of secondary recipient mice that were transplanted with BM cells of primary engrafted mice (BB n=13, CRLF2-IL7RAwt n=3, CRLF2-IL7RAins n=8, IL7RAwt-GFP n=9, IL7RAins-GFP n=10). Dot plots depict sample scatter with mean +/− SEM. Gray linkers indicate statistically significant difference (p < 0.05) between groups. Statistical analyses were performed using Kruskal-Wallis non-parametric one-way ANOVA test with Dunn’s post-hoc analysis.

One of the hallmarks of leukemic cells is the capability of self-renewal, a property of stem cells. To test whether expression of activated-IL7RA affects self-renewal we re-transplanted 100,000-150,000 transduced cells that were harvested from the BM of primary mice 28-32 weeks after transplantation. As portrayed in figure 3C, the capacity of IL7RA transduced cells harvested from primary mice to repopulate secondary mice was significantly enhanced when compared to cells from mice that were transplanted with BB transduced control. This was particularly evident in the activated-IL7RAins transduced group (p=0.0073, one-way ANOVA test).

### Initiation of de-novo leukemia after serial transplantation of activated-IL7RA hematopoietic progenitors

BCP-ALL is the end-result of sequential cumulative mutational events in which an initiation mutation causing “pre-leukemia” is followed by secondary mutations mediating progression to overt malignancy^3^. Transformation from pre-leukemia to leukemia in children is rare and often associated with an intervening period of several years. In agreement with this, none of the primary recipients of the transduced human hematopoietic progenitors developed leukemia within the first half year of follow up. This notwithstanding, we hypothesized that a selective pressure of serial transplantation might promote the evolution of pre-leukemic cells into leukemia.

Indeed, as depicted in figure 4, one of the five (two samples originated in the same batch) IL7Rains transduced cord bloods that developed a clear CD10^high^CD19^low^ population in the first transplanted mouse (bottom right sample in figure 3B) progressed to leukemia in a secondary transplanted mouse. This leukemia was characterized by expansion of CD34^+^CD10^+^CD19^+^ population (figure 4 A,B). The cells densely populated the BM and spleen of the engrafted mouse (supplementary figure 4). To validate that the human engrafted cells represented an overt leukemia, tertiary transplants were performed, in which all (nine out of nine) the recipients developed identical leukemia within 8-15 weeks of transplantation (supplementary figure 4).

**Figure 4.**
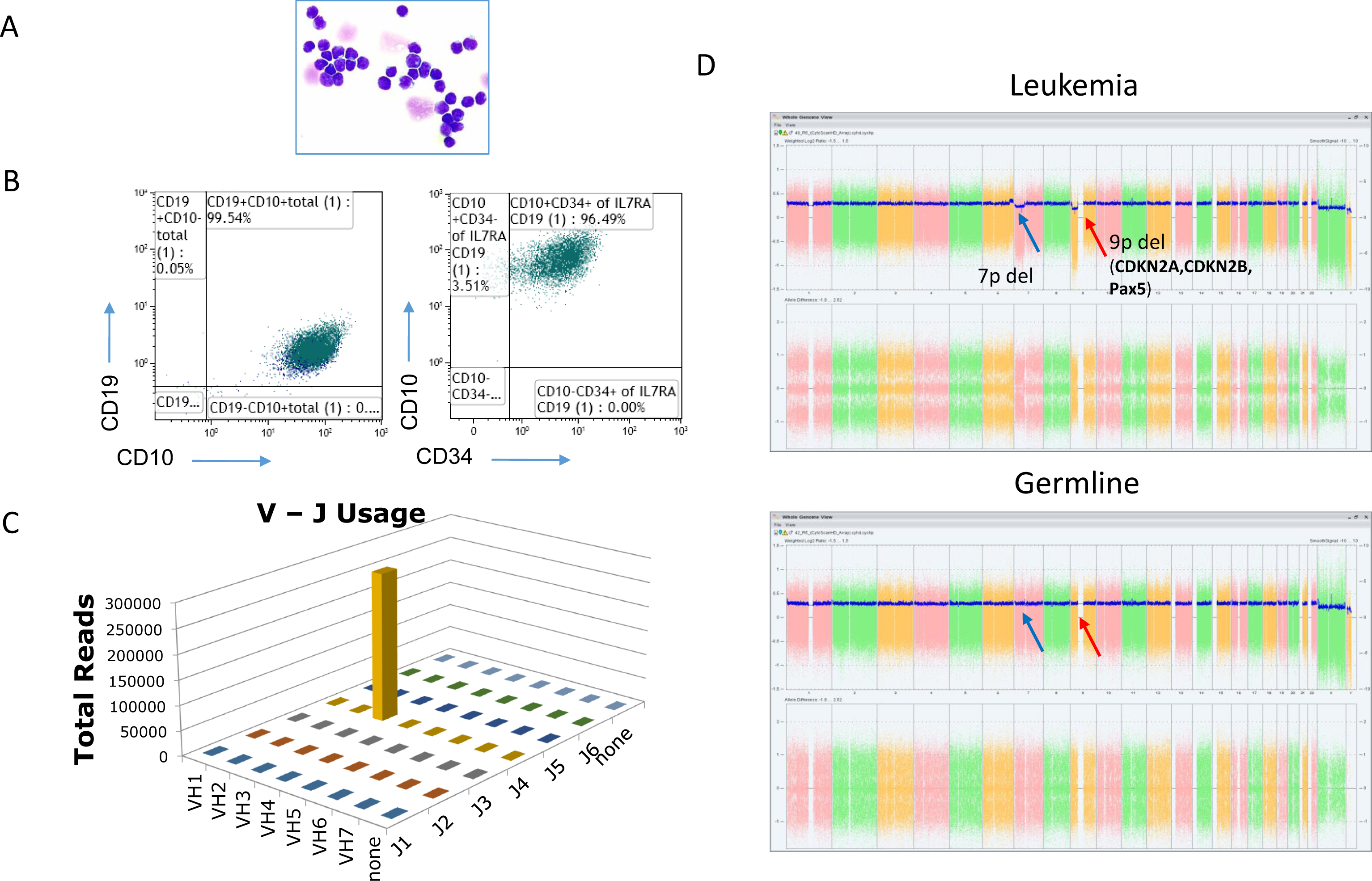
Secondary transplantation of IL7RA activated human hematopoietic progenitors result in development of clonal B-cell precursor leukemia. (A) GIEMSA staining of cytospin from BM of the leukemic mouse. (B) Flow cytometer scatter plot of human engrafted cells in BM of leukemic mouse. (C) Bar-graph of V-J rearrangements in leukemic population. The bars represent counts in the sequenced library of B-cell receptor rearrangements (D) SNP analysis of leukemic cells (Leukemia) and of BB transduced engrafted cells from the corresponding cord blood (representing germline).

VH-region sequencing of genomic DNA revealed that the leukemia was clonal and carried a non-functional (containing a stop codon) V3-15J4 gene rearrangement (figure 4C and supplementary figure 5). Only this one allele was non-productively rearranged in the leukemic cells in agreement with a block of differentiation in an early B-cell stage. This was further supported by mass cytometry analysis classifying the leukemic cells as pro-B-II population (supplementary figure 6).

**Figure 5.**
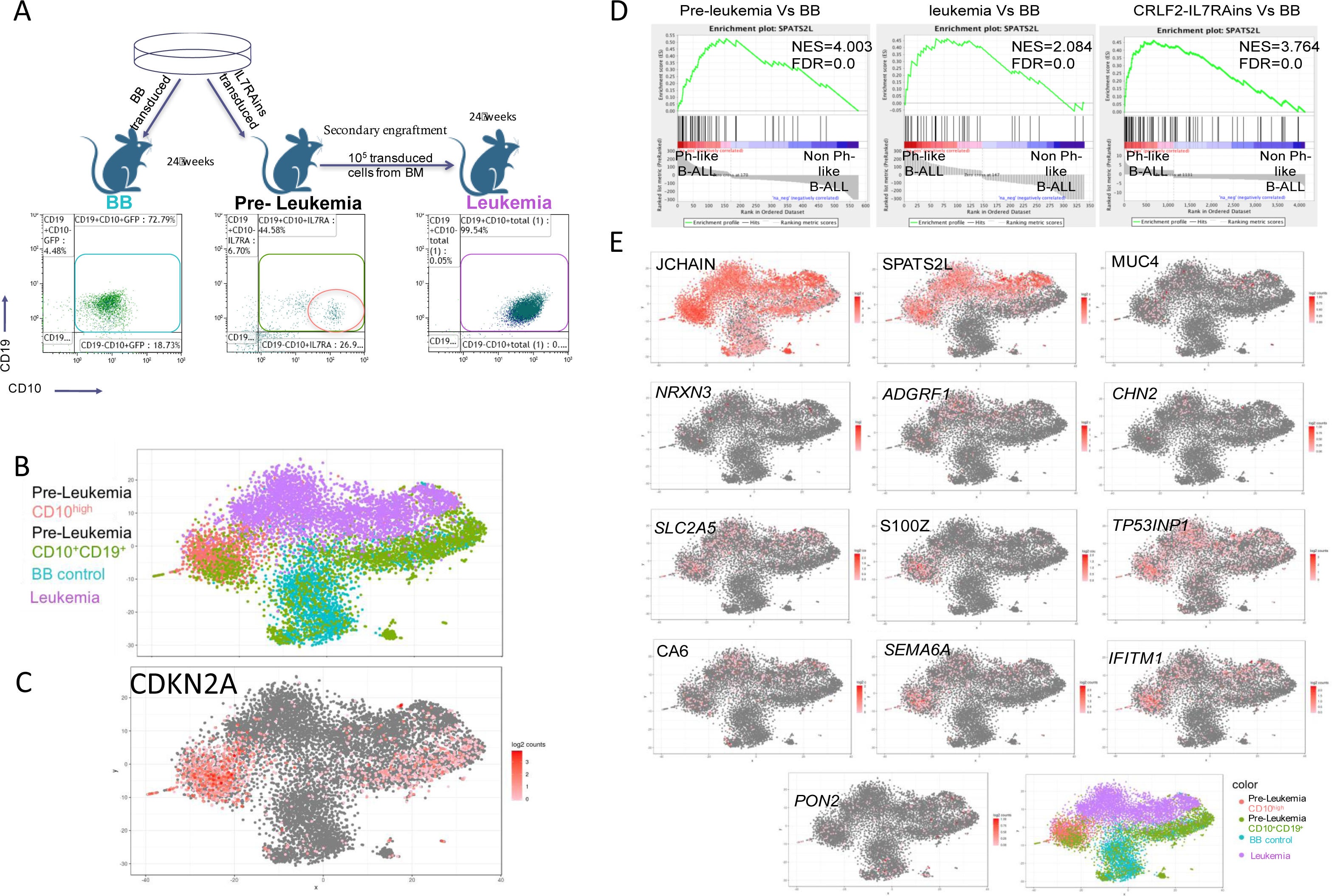
Philadelphia-like ALL gene signature in bulk and scRNAseq analyses of activated-IL7RA engrafted cells. (A) Scheme of sample acquisition for scRNAseq. (B)Transcriptome correlation t-SNE map after 10X scRNAseq. (C) Relative expression of CDKN2A displayed on t-SNE map. (D) GSEA enrichment plot of Pre-leukemic cells, leukemic cells and CRLF2-IL7RAins over BB differentially expressed genes aligned to Ph-like vs non Ph-like ranked gene list. (E) Relative expression of 13 genes that were detectable by scRNAseq out of 15 ph-like diagnosis clinical panel, displayed on t-SNE map.

Of note, repertoire sequencing of CD45^+^CD19^+^CD10^+^ cells from the primary mouse that generated the leukemia detected the leukemic clone at a frequency of 0.08%-0.02% of the CD10^high^CD19^low^ and CD10^med^CD19^+^ populations respectively (supplementary figure 7). Thus, the leukemia was derived from a pre-leukemic clone in the primary transplanted mouse. Karyotypic analysis and Cytoscan HD DNA array of the leukemic cells revealed several DNA copy number abnormalities (figure 4D and supplementary figure 8). Of special interest is the deletion in 9p13 that involves CDKN2A and PAX5 both of which are well characterized progression events in the evolution of BCP-ALL^36^. Genomic analysis also detected an internal IKZF1 deletion (supplementary figure 9), another typical progression event in high risk Philadelphia and Ph-like BCP-ALL^37–39^. Importantly, none of these abnormalities were detected in cells from the same CB transduced with the control backbone vector, confirming that they did not exist in the germline prior to the transduction with activated-IL7RAins receptor (figure 4D and supplementary figure 9). The co-existence of PAX5, CDKN2A and IKZF1 mutations classifies this leukemia as “Ikaros plus” high risk ALL^40^. Additionally, pSTAT5 analysis by flow cytometry demonstrated a Ph-like typical high basal activation of the JAK-STAT pathway^41^ that was cytokine independent (supplementary figure 10).

To better map the genetic landscape of the malignancy, we performed whole-genome sequencing (WGS, physical depth of 60×) of leukemia sample and matched engrafted cells from the same CB transduced with BB. As portrayed in supplementary table 3, seven genomic deletions outside the immunoglobulin loci were evident in the leukemic cells. This finding agrees with what was previously reported for *ETV6-RUNX1* human ALL^42^.

Leukemia developed only in one CB out of 13 batches of CB that were transduced with IL7RAins vector. Recent gene wide association (GWAS) studies have described several germline single nucleotide polymorphic (SNP) alleles thought to predispose to BCP-ALL^43–52^. Interestingly the original CB from which the overt leukemia was developed, carried five germline heterozygous risk SNPs in four genes. Of most interest are the GATA3 SNP (rs3824662) which was reported to closely associates with Ph-like B-ALL ^45, 47, 52^ and the CEBPE SNP previously predicted to reduce Ikaros binding to the CEBPE promoter and thus allow for elevated expression of CEBPE^49^ (rs2239635). Additional SNPs were detected in ARID5B locus (rs7089424^43^ and rs10821936^48, 51^), and CDKN2BAS locus rs564398^46^ (supplementary table 4). Although each of these alleles is relatively common the co-occurrence of five heterozygous SNPs is remarkable and, like humans, could potentially explain the propensity to transformation.

### Single cell analysis of B-cell precursor cells transduced with activated-IL7RA reveals a pre-leukemic population with a strong Ph-like gene signature

To better characterize the genetic changes that preceded full leukemic transformation we aimed to identify distinctive expression patterns unique for the leukemic and pre-leukemic cells. We hence sorted human CD10^+^CD19^+^ cells from the BM of the leukemic mouse, the pre-leukemic mouse (“pre-leukemia CD10^+^CD19^+^”) and the mouse engrafted with matching CB transduced with control BB virus. Additionally, we separately sorted the CD10^high^CD19^low^ sub-population presumably enriched with pre-leukemic cells because of their high resemblance to the leukemic immunophenotype (“pre-leukemia CD10^high^”). Cells were subjected to 10X single cell gene expression analysis (see experiment illustration figure 5A).

T-distributed somatic neighbor embedding (TSNE) plot demonstrated a clear separation between three cell populations: the leukemic cells, the BB control cells and the pre-leukemic CD10^high^ cells (*figure 5*B). The total CD10^+^CD19^+^ population of the pre-leukemic mouse was distributed between the CD10^high^ cluster and the BB control cluster.

Bulk differential expression analysis of the above populations shows a close hierarchical relationship between the total CD10^+^CD19^+^ from the pre-leukemic cells and the control backbone transduced cells (only 71 differentially expressed genes-supplementary figure 11A). In contrast, comparison of the leukemic cells and the CD10^high^ pre-leukemic cells to the BB transduced cells revealed 343 and 592 differentially expressed genes respectively (supplementary figure 11A). 166 of these differentially expressed genes were shared between the leukemia and pre-leukemia CD10^high^ groups, thus supporting the hypothesis that the CD10^high^CD19^low^ compartment is enriched with “pre-leukemic” cells.

Strikingly, CDKN2A expression which was significantly up-regulated in the IL7RA activated pre-leukemic CD10^high^ cluster (as was also in the bulk RNAseq analysis of CRLF2-IL7RAins – see supplementary table 1) was absent in the leukemic cluster (*figure 5*C) in which CDKN2A was deleted (figure *4*D). This observation strongly implies that the loss of the negative cell cycle regulator CDKN2A may be important in the leukemic evolution of IL7Rains pre-leukemic cells. Similar to CRLF2-IL7RAins driven RAG1/2 over expression observed in the bulk RNAseq data (supplementary table 1), elevated levels of RAG1 transcript were detected both in the pre-leukemic and in the leukemic populations (supplementary figure 11B). we speculate that similar to what was recently described during V(D)J recombination^53^ and specifically reported for ETV6-RUNX1 ALL^42^, increased RAG1/2 activity in B-cell precursor pre-leukemia might lead to genetic instability.

To investigate how closely the experimental leukemia recapitulated primary human Ph-like ALLs, we compiled ranked lists of the differentially expressed genes between the leukemia and the pre-leukemia CD10^high^ cells and control BB cells and used it in a gene set enrichment analysis against list of differentially expressed genes from two groups of BCP-ALL: Philadelphia and Ph-like BCP-ALL versus combined groups of BCP-ALL leukemias (patient database St. Jude’s group-GSE26281). As seen in *figure 5*D, a Philadelphia-like gene signature was found both in the leukemic and pre-leukemic sample when compared to the BB control (see major genes contributing to enrichment in supplementary table 5). Similar results were obtained from analysis of bulk RNAseq of CRLF2-IL7RAins vs BB transduced transplanted cells (*figure 5*D). Furthermore, to test the clinical relevance of our experimental model we assessed the expression of 15 genes that are clinically used in diagnosis of Ph-Like patients ^54^. Strikingly, 12 genes from this list out of 13 that were detected in the RNAseq analysis, were predominantly expressed in the pre-leukemic CD10^high^ and in the leukemic cells whereas only one (*PON2)*, did not show preferential expression pattern (*figure 5*E).

## Discussion

Childhood ALL is preceded by a clinically silent phase of pre-leukemia detectable only through molecular genomic approaches. While the pre-leukemia state is estimated to be fairly common (up to 1:20 children^55^) transformation to leukemia is rare. This observation places emphasis on understanding how the pre-leukemic phase is initiated and how the responsible lesions can pre-dispose cells for subsequent frank transformation. Experimental models that authentically re-capitulate disease initiation and progression in relevant human cells are thus required but are currently largely lacking.

Herein, we explore these issues in the most common subtype of the poor-prognosis Philadelphia-like (Ph-like) B-cell precursor acute lymphoblastic leukemia (BCP-ALL). This variant is commonly associated with aberrant expression of cytokine receptor-like factor-2 (CRLF2) which dimerizes with Interleukin-7 receptor alpha (IL7RA) to form the receptor for Thymic Stromal Lymphopoietin (TSLP). Additional mutations in IL7RA itself occurring with or without the aberrant expression of CRLF2, or its downstream signaling components JAK1 or JAK2, further underscore the importance of the IL7RA receptor axis in Ph-Like BCP-ALL^14, 15^. Whether IL7RA signaling has an instructive role in initiation of human BCP-ALL and if so how is the receptor activation driven pre-leukemia is predisposed to transformation, is unknown.

Here we provide the first experimental evidence in human hematopoietic cells, that expression of activated-IL7RA (IL7RAins), with or without CRLF2, modify human B-cell development into a state that could evolve to overt “Ph-like” BCP-ALL.

This “pre-leukemic” compartment of B-cell progenitors is enriched in early B-cell precursors with typical immunophenotype of BCP-ALL (CD34^+^CD10^+^CD19^+^). These cells carried an increased frequency of loci with only DJ and/or non-productive V(D)J rearrangements of the IgH chain. The survival of these cells, normally subject to apoptosis due to lack of antigenic selection, might be explained by the increased expression of BCL2L1, previously described to rescue pro-B cells with aberrant V(D)J rearrangements from apoptosis^56^. The expression of other BCL2 family members and pro-survival genes was also increased. Molecularly, these cells displayed the typical “Ph-like” leukemia gene expression signature and, interestingly, increased expression of the cell cycle inhibitor CDKN2A, possibly acting as a “gatekeeper” in restraining aberrant proliferation. This regulator was subsequently lost when these cells further evolved towards overt leukemia. We also observed significantly elevated expression of RAG1/2 in the transduced cells (supplementary table 1 and supplementary figure 11). This is not unexpected for B-cells that are held in an early progenitor stage and leads us to speculate that genetic instability driven by prolonged RAG activity might, as previously demonstrated ^42, 53, 57^, be involved in the eventual development of leukemia, for example by enabling deletions of CDKN2A and IKZF1.

A possible explanation to the early block of differentiation observed as a result of constant activation of IL7RA lays in the role played by IL7 signaling as was shown in normal mouse B-cell progenitor development; whereby down regulation of the pathway and switch to pre-BCR signaling, the expression of Bcl6 and elevation of Ikaros promotes normal progression of B-cell differentiation^58, 59^. Thus, our observations in human B-cell progenitors resonate well with the earlier finding that in mouse cells constitutive activation of IL7 signaling results in cell differentiation arrest at an early B-cell stage ^60^. This result demonstrates that IL7 role in human and mouse B-cell differentiation may be more similar than previously contemplated. Transduction of hematopoietic progenitors with IL7RAins led to the development of a population with enhanced self-renewal capacity as manifested by engraftment in secondary mouse recipients and the eventual development of leukemia in a single cord blood. Single cell RNAseq analysis (*figure 5*) suggested that pre-leukemic cells resided within a subpopulation of early B-cell precursors with CD34^+^CD10^high^CD19^low^ immunophenotype^61^. This population harbored the specific leukemic V(D)J clone. The experimental leukemia presented with all the hallmarks of Ph-like “IKZF1 plus” human leukemia including a Ph-like expression signature and the spontaneous acquisition of the genomic loss of IKZF1 and both alleles of the cell cycle regulator CDKN2A^16, 40, 62^ as well as PAX5.

The singularity of the leukemic event may not be surprising since rare secondary spontaneous genomic events are pre-requisite for progression of human pre-leukemia to leukemia. It has been estimated that leukemia develops in only 1% or less of children born with pre-leukemic clones^1, 55, 63, 64^. The time in children for development of leukemia is between 2-15 years while our experiment lasted only up to a year. Interestingly in children the CRLF2/IL7RA subtype occurs later than the other genotypic subtypes of ALL^65^. Recently, gene-wide association studies have discovered several relatively common SNPs conferring a higher risk for childhood ALL^45, 47, 52, 66–68^. Strikingly, the genotype of the specific CB the transduction of which resulted in BCP-ALL, revealed heterozygosity for five predisposing SNPs scattered across four of these genes. One is GATA3 which has previously been implicated in increased risk of Ph-like ALL^47, 52, 69^. Thus, the leukemia may have arisen in a CB (“a donor”) with a higher pre-disposition for BCP-ALL.

We designed this study to elucidate the role of the TSLP and IL7RA receptor pathway activation in the development of BCP-ALL. CRLF2 and IL7RA sequence and function differ between mouse and human^70–75^. Hence, in this work we studied human cells. The inability of sole expression of CRLF2 to produce a marked phenotype may be explained by the lack of cross reactivity of mouse Tslp with the human CRLF2 receptor as was demonstrated by Payne et al.^76^ The most common paired mutation with CRLF2 in BCP-ALL is activated JAK2R683^32, 77^. However, enforced expression of CRLF2-JAK2R683G in our experimental system was highly toxic to the transduced cells most likely due to hyper-activation of the JAK-STAT signaling pathway as we and others have described ^7, 78, 79^. Activating mutations in IL7RA were identified by us and others in both BCP and T ALLs^13, 15, 80, 81^. The most common mutations consist of in-frame insertion of cysteine (e.g the in-frame insertion PPCL used in this study) into the extracellular domain causing homodimerization of IL7RA, ligand independent activation and signaling through JAK1-STAT5. Thus, the model generated in our research is highly relevant to human leukemias. To properly study the role of human CRLF2, a mouse strain transgenic for human TSLP is currently being generated.

In summary, we demonstrate here that activation of IL7RA has an instructive role in the development of human B-cell precursor leukemia. It initiates a pre-leukemic state that can evolve to leukemia that recapitulates the natural Ph-like leukemia. As approaches for targeting the CRLF2 and/or IL7RA are currently under development these observations may bear therapeutic significance.

## Acknowledgements

We thank Michael Gershovits from the Mantoux bioinformatics institute of the Nancy and Stephen Grand Israel National Center for Personalized Medicine, Weizmann institute of science, for RNAseq analysis service, Dvir Dahary for help with WGS data analysis and Idit Shiff from the genomic applications laboratory, the core research facility, faculty of medicine – Ein Kerem, the Hebrew university of Jerusalem, Israel, for 10X RNA sequencing services. We are indebted to Nava Gershman and Itzhak Ben Moshe (Ofer) for help with xenograft experiments. We thank all past and present members of S.I. research group for fruitful discussions and advice.

We thank the Rawlings lab for sharing the Eu-B29 lentiviral construct.

This work was supported by the Israel Science Foundation Legacy and ICORE programs, Children with Cancer (UK), Swiss Bridge Foundation, WLBH Foundation, Waxman Cancer Research Foundation, US–Israel Binational Science Foundation, Israeli health ministry ERA-NET program, Hans Neufeld Stiftung, and Israel Cancer Research Foundation including ICRF-City of Hope Miller foundation. I.G was partially supported by Israeli ministry of Immigrant Absorption.

This work was performed in partial fulfilment of the requirements for a PhD degree of Ifat Geron, Sackler Faculty of Medicine, Tel Aviv University, Israel and Division of Biological Sciences University of California San Diego USA.

## Authorship Contributions

I.G., A.M.S., S.I. designed the study. I.G. and A.M.S. performed most of the experiments. N.T preformed initial IL7RA experiments. I.G. A.M.S., J.B. V.T. C.J., A.R.G performed and analysed WGS and/or bulk RNAseq experiments. I.R.K. interpreted Immunoseq V(D)J rearrangements analysis. I.G. A.M.S., O.P. performed and interpreted 10XscRNAseq. U.F., A.B., preformed and interpreted exome sequencing. J.S and K.L.D preformed and analysed mass cytometry experiments. Y.N.L performed VH-region sequencing of Leukemic cells. V.M. performed Karyotype analysis of the leukemic cells M.H. performed SNP analysis. I.M and H.F provided technical support for experiments. Y.B. and M.M critically reviewed the experiments and provided important advice. A.N. provided cord blood samples and critically reviewed experiments. I.G., A.M.S., T.E. and S.I. analyzed and interpreted the data. I.G. and S.I. wrote the manuscript.

## Disclosure of Conflicts of Interest

IRK is a full time employee of Adaptive Biotechnologies, Inc.

## References

1. Greaves MF, Wiemels J. Origins of chromosome translocations in childhood leukaemia. Nat Rev Cancer. 2003;3(9):639–649.

2. Gu Z, Churchman ML, Roberts KG, et al. PAX5-driven subtypes of B-progenitor acute lymphoblastic leukemia. Nature Genetics. 2019.

3. Hong D, Gupta R, Ancliff P, et al. Initiating and cancer-propagating cells in TEL-AML1-associated childhood leukemia. Science. 2008;319(5861):336–339.

4. Oshima K, Khiabanian H, da Silva-Almeida AC, et al. Mutational landscape, clonal evolution patterns, and role of RAS mutations in relapsed acute lymphoblastic leukemia. Proc Natl Acad Sci U S A. 2016;113(40):11306–11311.

5. Malinowska-Ozdowy K, Frech C, Schonegger A, et al. KRAS and CREBBP mutations: a relapse-linked malicious liaison in childhood high hyperdiploid acute lymphoblastic leukemia. Leukemia. 2015;29(8):1656–1667.

6. Irving J, Matheson E, Minto L, et al. Ras pathway mutations are prevalent in relapsed childhood acute lymphoblastic leukemia and confer sensitivity to MEK inhibition. Blood. 2014;124(23):3420–3430.

7. Schwartzman O, Savino AM, Gombert M, et al. Suppressors and activators of JAK-STAT signaling at diagnosis and relapse of acute lymphoblastic leukemia in Down syndrome. Proc Natl Acad Sci U S A. 2017;114(20):E4030–E4039.

8. Puel A, Ziegler SF, Buckley RH, Leonard WJ. Defective IL7R expression in T(-)B(+)NK(+) severe combined immunodeficiency. Nat Genet. 1998;20(4):394–397.

9. Roifman CM, Zhang J, Chitayat D, Sharfe N. A partial deficiency of interleukin-7R alpha is sufficient to abrogate T-cell development and cause severe combined immunodeficiency. Blood. 2000;96(8):2803–2807.

10. Parrish YK, Baez I, Milford T-A, et al. IL-7 Dependence in Human B Lymphopoiesis Increases During Progression of Ontogeny from Cord Blood to Bone Marrow. Journal of immunology (Baltimore, Md : 1950). 2009;182(7):4255–4266.

11. Prieyl JA, LeBien TW. Interleukin 7 independent development of human B cells. Proc Natl Acad Sci U S A. 1996;93(19):10348–10353.

12. Barata JT, Durum SK, Seddon B. Flip the coin: IL-7 and IL-7R in health and disease. Nature Immunology. 2019;20(12):1584–1593.

13. Zenatti PP, Ribeiro D, Li W, et al. Oncogenic IL7R gain-of-function mutations in childhood T-cell acute lymphoblastic leukemia. Nat Genet. 2011;43(10):932–939.

14. Shochat C, Tal N, Bandapalli OR, et al. Gain-of-function mutations in interleukin-7 receptor-alpha (IL7R) in childhood acute lymphoblastic leukemias. The Journal of experimental medicine. 2011;208(5):901–908.

15. Reshmi SC, Harvey RC, Roberts KG, et al. Targetable kinase gene fusions in high-risk B-ALL: a study from the Children’s Oncology Group. Blood. 2017;129(25):3352–3361.

16. Roberts KG, Li Y, Payne-Turner D, et al. Targetable kinase-activating lesions in Ph-like acute lymphoblastic leukemia. N Engl J Med. 2014;371(11):1005–1015.

17. Den Boer ML, van Slegtenhorst M, De Menezes RX, et al. A subtype of childhood acute lymphoblastic leukaemia with poor treatment outcome: a genome-wide classification study. Lancet Oncol. 2009;10(2):125–134.

18. Izraeli S. Beyond Philadelphia: ‘Ph-like’ B cell precursor acute lymphoblastic leukemias – diagnostic challenges and therapeutic promises. Curr Opin Hematol. 2014;21(4):289–296.

19. Potter N, Jones L, Blair H, et al. Single-cell analysis identifies CRLF2 rearrangements as both early and late events in Down syndrome and non-Down syndrome acute lymphoblastic leukaemia. Leukemia. 2019;33(4):893–904.

20. Morak M, Attarbaschi A, Fischer S, et al. Small sizes and indolent evolutionary dynamics challenge the potential role of P2RY8-CRLF2-harboring clones as main relapse-driving force in childhood ALL. Blood. 2012;120(26):5134–5142.

21. Vesely C, Frech C, Eckert C, et al. Genomic and transcriptional landscape of P2RY8-CRLF2-positive childhood acute lymphoblastic leukemia. Leukemia. 2016.

22. Lane AA, Chapuy B, Lin CY, et al. Triplication of a 21q22 region contributes to B cell transformation through HMGN1 overexpression and loss of histone H3 Lys27 trimethylation. Nat Genet. 2014;46(6):618–623.

23. Russell LJ, Capasso M, Vater I, et al. Deregulated expression of cytokine receptor gene, CRLF2, is involved in lymphoid transformation in B-cell precursor acute lymphoblastic leukemia. Blood. 2009;114(13):2688–2698.

24. Yokoyama K, Yokoyama N, Izawa K, et al. In vivo leukemogenic potential of an interleukin 7 receptor alpha chain mutant in hematopoietic stem and progenitor cells. Blood. 2013;122(26):4259–4263.

25. Sather BD, Ryu BY, Stirling BV, et al. Development of B-lineage predominant lentiviral vectors for use in genetic therapies for B cell disorders. Mol Ther. 2011;19(3):515–525.

26. Tiscornia G, Singer O, Verma IM. Production and purification of lentiviral vectors. Nat Protoc. 2006;1(1):241–245.

27. Carlson CS, Emerson RO, Sherwood AM, et al. Using synthetic templates to design an unbiased multiplex PCR assay. Nat Commun. 2013;4:2680.

28. Good Z, Sarno J, Jager A, et al. Single-cell developmental classification of B cell precursor acute lymphoblastic leukemia at diagnosis reveals predictors of relapse. Nat Med. 2018;24(4):474–483.

29. Lu L, Osmond DG. Apoptosis during B lymphopoiesis in mouse bone marrow. J Immunol. 1997;158(11):5136–5145.

30. Shinkai Y, Rathbun G, Lam KP, et al. RAG-2-deficient mice lack mature lymphocytes owing to inability to initiate V(D)J rearrangement. Cell. 1992;68(5):855–867.

31. Korsmeyer SJ, Arnold A, Bakhshi A, et al. Immunoglobulin gene rearrangement and cell surface antigen expression in acute lymphocytic leukemias of T cell and B cell precursor origins. J Clin Invest. 1983;71(2):301–313.

32. Hertzberg L, Vendramini E, Ganmore I, et al. Down syndrome acute lymphoblastic leukemia, a highly heterogeneous disease in which aberrant expression of CRLF2 is associated with mutated JAK2: a report from the International BFM Study Group. Blood. 2010;115(5):1006–1017.

33. Kharabi Masouleh B, Geng H, Hurtz C, et al. Mechanistic rationale for targeting the unfolded protein response in pre-B acute lymphoblastic leukemia. Proc Natl Acad Sci U S A. 2014;111(21):E2219–2228.

34. Ghia P, ten Boekel E, Sanz E, de la Hera A, Rolink A, Melchers F. Ordering of human bone marrow B lymphocyte precursors by single-cell polymerase chain reaction analyses of the rearrangement status of the immunoglobulin H and L chain gene loci. J Exp Med. 1996;184(6):2217–2229.

35. LeBien TW. Fates of human B-cell precursors. Blood. 2000;96(1):9–23.

36. Mullighan CG, Phillips LA, Su X, et al. Genomic analysis of the clonal origins of relapsed acute lymphoblastic leukemia. Science. 2008;322(5906):1377–1380.

37. Mullighan CG, Su X, Zhang J, et al. Deletion of IKZF1 and prognosis in acute lymphoblastic leukemia. The New England journal of medicine. 2009;360(5):470–480.

38. Joshi I, Yoshida T, Jena N, et al. Loss of Ikaros DNA-binding function confers integrin-dependent survival on pre-B cells and progression to acute lymphoblastic leukemia. Nat Immunol. 2014;15(3):294–304.

39. Mullighan CG, Miller CB, Radtke I, et al. BCR-ABL1 lymphoblastic leukaemia is characterized by the deletion of Ikaros. Nature. 2008;453(7191):110–114.

40. Stanulla M, Dagdan E, Zaliova M, et al. IKZF1plus Defines a New Minimal Residual Disease–Dependent Very-Poor Prognostic Profile in Pediatric B-Cell Precursor Acute Lymphoblastic Leukemia. Journal of Clinical Oncology. 2018;36(12):1240–1249.

41. Tasian SK, Doral MY, Borowitz MJ, et al. Aberrant STAT5 and PI3K/mTOR pathway signaling occurs in human CRLF2-rearranged B-precursor acute lymphoblastic leukemia. Blood. 2012;120(4):833–842.

42. Papaemmanuil E, Rapado I, Li Y, et al. RAG-mediated recombination is the predominant driver of oncogenic rearrangement in ETV6-RUNX1 acute lymphoblastic leukemia. Nat Genet. 2014;46(2):116–125.

43. Papaemmanuil E, Hosking FJ, Vijayakrishnan J, et al. Loci on 7p12.2, 10q21.2 and 14q11.2 are associated with risk of childhood acute lymphoblastic leukemia. Nat Genet. 2009;41(9):1006–1010.

44. Hungate EA, Vora SR, Gamazon ER, et al. A variant at 9p21.3 functionally implicates CDKN2B in paediatric B-cell precursor acute lymphoblastic leukaemia aetiology. Nat Commun. 2016;7:10635.

45. Perez-Andreu V, Roberts KG, Xu H, et al. A genome-wide association study of susceptibility to acute lymphoblastic leukemia in adolescents and young adults. Blood. 2015;125(4):680–686.

46. Iacobucci I, Sazzini M, Garagnani P, et al. A polymorphism in the chromosome 9p21 ANRIL locus is associated to Philadelphia positive acute lymphoblastic leukemia. Leuk Res. 2011;35(8):1052–1059.

47. Perez-Andreu V, Roberts KG, Harvey RC, et al. Inherited GATA3 variants are associated with Ph-like childhood acute lymphoblastic leukemia and risk of relapse. Nat Genet. 2013;45(12):1494–1498.

48. Yang W, Trevino LR, Yang JJ, et al. ARID5B SNP rs10821936 is associated with risk of childhood acute lymphoblastic leukemia in blacks and contributes to racial differences in leukemia incidence. Leukemia. 2010;24(4):894–896.

49. Wiemels JL, de Smith AJ, Xiao J, et al. A functional polymorphism in the CEBPE gene promoter influences acute lymphoblastic leukemia risk through interaction with the hematopoietic transcription factor Ikaros. Leukemia. 2016;30(5):1194–1197.

50. Pui CH, Roberts KG, Yang JJ, Mullighan CG. Philadelphia Chromosome-like Acute Lymphoblastic Leukemia. Clin Lymphoma Myeloma Leuk. 2017;17(8):464–470.

51. Trevino LR, Yang W, French D, et al. Germline genomic variants associated with childhood acute lymphoblastic leukemia. Nat Genet. 2009;41(9):1001–1005.

52. Migliorini G, Fiege B, Hosking FJ, et al. Variation at 10p12.2 and 10p14 influences risk of childhood B-cell acute lymphoblastic leukemia and phenotype. Blood. 2013;122(19):3298–3307.

53. Kirkham CM, Scott JNF, Wang X, et al. Cut-and-Run: A Distinct Mechanism by which V(D)J Recombination Causes Genome Instability. Mol Cell. 2019.

54. Kang H, Roberts KG, Chen I-ML, et al. Development and Validation Of a Highly Sensitive and Specific Gene Expression Classifier To Prospectively Screen and Identify B-Precursor Acute Lymphoblastic Leukemia (ALL) Patients With a Philadelphia Chromosome-Like (“<em>Ph-like</em>” or “<lt;em>BCR-ABL1-Like<lt;/em>”) Signature For Therapeutic Targeting and Clinical Intervention. Blood. 2013;122(21):826–826.

55. Schafer D, Olsen M, Lahnemann D, et al. Five percent of healthy newborns have an ETV6-RUNX1 fusion as revealed by DNA-based GIPFEL screening. Blood. 2018;131(7):821–826.

56. Fang W, Mueller DL, Pennell CA, et al. Frequent aberrant immunoglobulin gene rearrangements in pro-B cells revealed by a bcl-xL transgene. Immunity. 1996;4(3):291–299.

57. Kitagawa Y, Inoue K, Sasaki S, et al. Prevalent involvement of illegitimate V(D)J recombination in chromosome 9p21 deletions in lymphoid leukemia. J Biol Chem. 2002;277(48):46289–46297.

58. Heizmann B, Kastner P, Chan S. Ikaros is absolutely required for pre-B cell differentiation by attenuating IL-7 signals. J Exp Med. 2013;210(13):2823–2832.

59. Duy C, Yu JJ, Nahar R, et al. BCL6 is critical for the development of a diverse primary B cell repertoire. J Exp Med. 2010;207(6):1209–1221.

60. Purohit SJ, Stephan RP, Kim H-G, Herrin BR, Gartland L, Klug CA. Determination of lymphoid cell fate is dependent on the expression status of the IL-7 receptor. The EMBO Journal. 2003;22(20):5511–5521.

61. Bendall SC, Davis KL, Amir el AD, et al. Single-cell trajectory detection uncovers progression and regulatory coordination in human B cell development. Cell. 2014;157(3):714–725.

62. Mullighan CG, Su X, Zhang J, et al. Deletion of IKZF1 and prognosis in acute lymphoblastic leukemia. N Engl J Med. 2009;360(5):470–480.

63. Mori H, Colman SM, Xiao Z, et al. Chromosome translocations and covert leukemic clones are generated during normal fetal development. Proceedings of the National Academy of Sciences of the United States of America. 2002;99(12):8242–8247.

64. Greaves MF, Maia AT, Wiemels JL, Ford AM. Leukemia in twins: lessons in natural history. Blood. 2003;102(7):2321–2333.

65. Buitenkamp TD, Izraeli S, Zimmermann M, et al. Acute lymphoblastic leukemia in children with Down syndrome: a retrospective analysis from the Ponte di Legno study group. Blood. 2014;123(1):70–77.

66. Vijayakrishnan J, Kumar R, Henrion MY, et al. A genome-wide association study identifies risk loci for childhood acute lymphoblastic leukemia at 10q26.13 and 12q23.1. Leukemia. 2017;31(3):573–579.

67. Vijayakrishnan J, Studd J, Broderick P, et al. Genome-wide association study identifies susceptibility loci for B-cell childhood acute lymphoblastic leukemia. Nat Commun. 2018;9(1):1340.

68. Wiemels JL, Walsh KM, de Smith AJ, et al. GWAS in childhood acute lymphoblastic leukemia reveals novel genetic associations at chromosomes 17q12 and 8q24.21. Nat Commun. 2018;9(1):286.

69. Madzio J, Pastorczak A, Sedek L, et al. GATA3 germline variant is associated with CRLF2 expression and predicts outcome in pediatric B-cell precursor acute lymphoblastic leukemia. Genes, Chromosomes and Cancer. 2019.

70. Jiang Q, Li WQ, Aiello FB, et al. Cell biology of IL-7, a key lymphotrophin. Cytokine Growth Factor Rev. 2005;16(4-5):513–533.

71. Giliani S, Mori L, de Saint Basile G, et al. Interleukin-7 receptor alpha (IL-7Ralpha) deficiency: cellular and molecular bases. Analysis of clinical, immunological, and molecular features in 16 novel patients. Immunol Rev. 2005;203:110–126.

72. Mazzucchelli R, Durum SK. Interleukin-7 receptor expression: intelligent design. Nat Rev Immunol. 2007;7(2):144–154.

73. He R, Geha RS. Thymic stromal lymphopoietin. Annals of the New York Academy of Sciences. 2010;1183:13–24.

74. Reche PA, Soumelis V, Gorman DM, et al. Human thymic stromal lymphopoietin preferentially stimulates myeloid cells. J Immunol. 2001;167(1):336–343.

75. Tonozuka Y, Fujio K, Sugiyama T, Nosaka T, Hirai M, Kitamura T. Molecular cloning of a human novel type I cytokine receptor related to δ1/TSLPR. Cytogenetic and Genome Research. 2001;93(1-2):23–25.

76. Francis OL, Milford TM, Martinez SR, et al. A novel xenograft model to study the role of TSLP-induced CRLF2 signals in normal and malignant human B lymphopoiesis. Haematologica. 2015.

77. Bercovich D, Ganmore I, Scott LM, et al. Mutations of JAK2 in acute lymphoblastic leukaemias associated with Down’s syndrome. Lancet. 2008;372(9648):1484–1492.

78. Grundschober E, Hoelbl-Kovacic A, Bhagwat N, et al. Acceleration of Bcr-Abl+ leukemia induced by deletion of JAK2. Leukemia. 2014;28(9):1918–1922.

79. Shojaee S, Caeser R, Buchner M, et al. Erk Negative Feedback Control Enables Pre-B Cell Transformation and Represents a Therapeutic Target in Acute Lymphoblastic Leukemia. Cancer Cell. 2015;28(1):114–128.

80. Shochat C, Tal N, Bandapalli OR, et al. Gain-of-function mutations in interleukin-7 receptor-{alpha} (IL7R) in childhood acute lymphoblastic leukemias. J Exp Med. 2011;208(5):901–908.

81. Shochat C, Tal N, Gryshkova V, et al. Novel activating mutations lacking cysteine in type I cytokine receptors in acute lymphoblastic leukemia. Blood. 2014;124(1):106–110.

