## Supplemental material for "An instructive role for IL7RA in the development of human B-cell precursor leukemia"

#### Supplemental data

##### Supplementary Methods

###### CRLF2 and IL7RA cloning

Bi-cistronic cloning of CRLF2 and IL7RA (wild type and mutated PPCL ins) was performed as following: For first position (CRLF2-GFP/ IL7RAwt/ins-GFP) the genes were amplified (Phusion High-Fidelity PCR Master Mix (Finnzymes, Espoo, Finland)) from previously cloned cDNA [1,2] using the primers: CRLF2 (first) left 5'-atatgaattcgagggcatggggcggctggt-3 right 5'-aagcggccgccacaacgccacgta-3. IL7RA(first) left 5'- atatgaattcccaccatgacaattctagg-3' right 5'-atatgcggccgcctggttttggtagaagctgga-3'. The purified products were cloned into pCDH-EF1$\alpha$-MCS-T2A-copGFP vector (Mountain View, CA) in Ecor1I and Not1 sites. For second position (CRLF2-

IL7RAwt/ins) IL7RA was amplified using IL7RA (second) left 5'-ATATTCCGGAATGACAATTCTAGG 3' and IL7RA (second) right (5'-CAGCATGTCGACCTACTGGTTTTGGTAGAAGCTGGA-3') The purified products were cloned into pCDH-EF1α-CRLF2-T2A-copGFP in BspEI and SalI sites.

pRRL Eμ B29 GFP WPRE vector was kindly provided by Rawlings lab[3] the original GFP was excised and an additional restriction site (NheI) was added in the following way: The B29 promoter was re-amplified with forward primer: 5' TCGATGATACCCTGATGAAGC 3' and reverse primer carrying NheI and Kozak sequence in the 3' end

5' TATATGTCGACGCTAGCGGTGGCGGTCCACTGCTCTGTCTC 3'. This PCR product and pRRL-Eμ-B29-GFP-WPRE vector were digested with SalI and XcmI. Purified products were ligated.

The bi-cistronic cassettes were excised from pCDH vector with NheI and SalI. And inserted to the altered pRRL-Eμ-B29 vector.

**Virus production:**

Production of lenti vector was done as described in[4]. In short: 3rd generation lenti vector packaging plasmids were co transfected 293T cells in the ratio of (15:10:5:4) (Lenti vector:pMDL:pVSVG:pREV) using ProFection Calcium Phosphate mammalian transfection system (Promega) according to manufacturer's protocol. Transfection medium was replaced 6-15 hour after transfection with 5% serum DMEM serum and virus-containing supernatant was collected 24 and 48 hours after replacement. Supernatant was then filtered with 0.45um PVDF filters (Millipore, Massachusetts, USA) and centrifuged in ultra-centrifuge using SW28 rotor for two and a half hours in

19,400r.p.m (70,000g). The virus was reconstituted in 300-600 SFEM medium (STEMCELL technologies, Vancouver, British Columbia, Canada). Concentrated virus was frozen in -80$^0$C until use. An aliquot of frozen virus was used for titer in 018Z cells percentage of transduced cells was evaluated by flow cytometry using GFP, CRLF2 or IL7RA antibodies (Biolegend California, USA) Titer (infectious units/ml) was calculated according to the following equation:

$$\frac{\%transduced\ cellsX\#cells\ at\ day\ of\ transduction}{total\ \mu l\ of\ virus/well}X1000 = virus\ IU/ml$$

**Transduction of CB CD34$^+$ hematopoietic progenitors**

5x10$^4$ – 7.5x10$^4$ CB CD34$^+$ cells were plated in 96 U bottom-well plate (Corning Incorporated, NY, USA) in 50-100µl SFEM (STEMCELL Technologies Vancouver, British Columbia, Canada) supplemented with hSCF (100ng/ml) hFLT3 ligand (100ng/ml) TPO (20ng/µl) and IL-6 (20ng/µl). Cells are transduced twice in consecutive days by addition of virus in MOI of 50-200 and spin (800g 32$^0$C 45 min no break). 4-8 hours after spin the wells are supplemented with fresh media. Prior to the second transduction, old media containing virus is discarded. Transduction efficiency is evaluated by flow cytometry using GFP or CRLF2/IL7RA antibodies.

**Flow cytometry and sorting**

Standard staining protocols were used for sort and analysis of cells. In brief, cells were washed in staining media (2%FBS in PBS) and re-suspended in of staining media containing fluorochrome-conjugated antibodies, blocking antibodies when mouse tissue was used and 7AAD for 30 min. (Supplementary table 6). Following staining, cells were washed and analyzed on Gallios flow cytometer (Beckman-Coulter, California, USA) or sorted using ARIA I/Aria III FACS sorter (BD Biosciences, San Jose, CA USA). Single stains and FMOs (Full minus one staining) of each fluorophore were used for cytometer setup and gating. Analysis was preformed using Kaluza software (Beckman-Coulter, California, USA) on live cells after exclusion of 7AAD positive stained cells.

For xenografts sample analysis Hematopoietic tissues (Spleen, Bone marrow (BM) and liver) and peripheral blood (PB) were harvested from mice at sacrifice time and kept throughout the processing time on ice. BM cells were flashed from the hind leg bones and strained through a 70μm mesh cell strainer. Spleen and liver were mashed on a 70μm mesh cell strainer. PB and spleen were subjected to red blood cell lysis (Biolegend, San Diego, CA, USA) per manufacturer's protocol. Cells that were not used for analysis/sort were viably frozen in FBS+10% DMSO.

For RNAseq and repertoire analysis, processed xenograft samples were stained as described above. For RNAseq, 5000-20000 Live $CD45^+$ $CD3^-$ CRLF2/$GFP^+$ cells were sorted directly into mini-centrifuge tubes containing 800μl cold TRIzol (ThermoFisher Scientific Waltham, MA USA) Tubes were vortexed immediately after sort and flash frozen in liquid nitrogen for further RNA purification. For repertoire analysis, 5000-20000 Live $CD45^+$ CRLF2/$GFP^+$ $CD10^+$ and $CD19^+$ cells were sorted directly into mini-centrifuge tubes containing 200ul STM. Cells were then pelleted at 800RPM for 10 minutes and kept at -20 or processed immediately for gDNA extraction. For single cell RNAseq 4000-10000 cells were sorted

For phosphorylation assays, cells (from sub confluent culture) were first washed and starved for four hours (in media with no cytokines). Cells were then incubated with cytokines (hIL7, hTSLP) for 20 minutes, washed and stained with LIVE/DEAD Fixable staining antibody per manufacturer's protocol [Thermo Fisher Scientific Waltham, MA

USA (molecular probes brand)], cells were then stained for cell surface markers, fixed with 1.5% formaldehyde for 10 minutes, porated with ice-cold MeOH while vigorously vortexing and incubated at 4°C for at least 10 min. cells were then stored over night or more (up to 2 weeks) in -20. Fixed cells were then washed twice in staining media then resuspended in staining media containing pSTAT antibodies and re-stained for surface markers. Stained cells were analyzed on Gallios™ Flow Cytometer (Beckman-Coulter, California, USA).

*Mass cytometry analysis*

Samples were processed as previously described (ref). Briefly, bone marrow samples (backbone (n=3), CRLF2/IL7ins (n=3), IL7ins (n=4), and healthy BM (n=3)) were thawed, stained with cisplatin to determine viability, rested for 30 minutes at 37°C and then perturbed with IL-7 (100 ng/mL) for 15 minutes (only for AAF49A and BM 22) before being fixed with formaldehyde 1.6% for 10 minutes at room temperature. Cells were then barcoded using palladium-based labeling reagents, collected in one tube, stained with surface antibodies and after being permeabilized with methanol stained with intracellular antibodies (

| **Protein** | **Clone** | **Manufacturer** | **Metal Isotope** | **Staining** |
|---|---|---|---|---|
| 4EBP1(pT36/T46) | 236B4 | Cell Signaling Technology | Nd144 | Intracellular |
| Akt (pS473) | D9E | Cell Signaling Technology | Tb159 | Intracellular |
| BTK (pY551/511) | 24A/BTK | BD Biosciences | Yb174 | Intracellular |
| cCaspase3 | C92-605 | BD Biosciences | Ho165 | Intracellular |
| CD10 | HI10a | Biolegend | Gd156 | Surface |
| CD127 | A019D5 | Biolegend | Dy162 | Surface |
| CD16 | 3G8 | Fluidigm | Bi209 | Surface |
| CD179a | HSL96 | Biolegend | Sm149 | Intracellular |
| CD179b | HSL11 | Biolegend | Gd158 | Intracellular |
| CD19 | H1B19 | Biolegend | Nd142 | Surface |
| CD20 | 2H7 | Biolegend | Sm147 | Surface |
| CD22 | HIB22 | Biolegend | Nd143 | Surface |
| CD235 | HIR2 | Biolegend | In115 | Surface |
| CD24 | ML5 | Biolegend | Gd160 | Surface |
| CD3 | UCHT1 | Biolegend | Er170 | Surface |
| CD34 | 581 | Biolegend | Nd148 | Surface |
| CD38 | HIT2 | Biolegend | Er168 | Surface |
| CD43 | CD43-10G7 | Biolegend | Er167 | Surface |
| CD45 human | HI30 | Fluidigm | Y89 | Surface |
| CD45 mouse | 30F11 | Biolegend | In113 | Surface |
| CD79b | CB3-1 | Biolegend | Nd146 | Surface |
| cPARP | F21-852 | BD Biosciences | La139 | Intracellular |
| Creb (pS133) | 87G3 | Cell Signaling Technology | Yb176 | Intracellular |
| CRLF2 | 1A6 | eBioscience | Dy161 | Surface |

| | | | | |
|---|---|---|---|---|
| CyclinA (total) | BF-683 | BD Biosciences | Sm154 | Intracellular |
| CyclinB1 (total) | GNS-1 | BD Biosciences | Dy164 | Intracellular |
| Erk1/2 (pT202/pY204) | D13-14-4E | Cell Signaling Technology | Yb173 | Intracellular |
| Glucocorticoid Receptor | D8H2 | Cell Signaling Technology | Eu151 | Intracellular |
| GFP | SF12.4 | Fluidigm | Tm169 | Intracellular |
| HistoneH3 (pS28) | HTA28 | Biolegend | Ce140 | Intracellular |
| IgHintracellular | polyclonal | Novus | Eu153 | Intracellular |
| IgH surface | MHM-98 | Fluidigm | Yb172 | Surface |
| Ikaros (total) | D10E5 | Cell Signaling Technology | Nd145 | Intracellular |
| Ki67 | B56 | BD Biosciences | Sm152 | Intracellular |
| PU.1 | 9G7 | Cell Signaling Technology | Gd157 | Intracellular |
| RB (pS807/811) | J112-906 | BD Biosciences | Er166 | Intracellular |
| rpS6 (pS235/pS236) | N7-548 | BD Biosciences | Lu175 | Intracellular |
| SRC (pY418) | K98-37 | BD Biosciences | Pr141 | Intracellular |
| STAT5 (pY694) | 47 | BD Biosciences | Gd155 | Intracellular |
| Syk (pY319/pY352) | 17a | BD Biosciences | Yb171 | Intracellular |
| TdT | E17-1519 | BD Biosciences | Dy163 | Intracellular |

Supplementary table 7). Finally cells were stained with 191/193Ir DNA intercalator before being analyzed the Helios mass cytometer (Fluidigm, Inc., South San Francisco, CA). Normalization of signal intensity loss during the CyTOF run was controlled utilizing metal standard beads mixed with the sample during the data acquisition.

Mass cytometry data were then analyzed using Cytobank (Cytobank Inc. Mountain View, CA) and were run through a B-cell developmental classifier recently described (Good Z Nat Med 2018[5] ).

Specifically, healthy bone marrow (run with the samples) was manually gated into 11 consecutive developmental stages of B-lymphopoiesis. The mean arsinh-transformed expression of 10 markers (CD45, CD20, CD24, CD34, CD38, IgMi, TdT, CD19, IgMs, CD10) was determined for each healthy population and single cells from each sample were assigned to the most similar healthy population based on the shorted Mahalanobis distance calculated from expression of the same 10 markers.

#### RNA/DNA sequencing and Expression profile analysis

*Bulk RNA Differential expression analysis*

Paired expression analysis of CRLF2-IL7RAins versus BB was performed as following: The $Log_2$ of the counts+1 was first calculated. The fold change (FC) of expression was defined as the differences between $Log_2$ CRLF2-IL7RAins counts and $Log_2$BB counts within the same cord blood batch. Significantly differential expressed genes were ranked per the average FC or their paired t values calculated as $t = \frac{\bar{d}-0}{S_d/\sqrt{n}}$ when $\bar{d}$ =avarage FC, n=6 pairs of samples and $S_d = \sqrt{\frac{1}{n-1}\{(\sum_1^n d_i^2) - n\bar{d}^2\}}$. Genes with overall low counts (background levels) in both samples (CRLF2IL7RAins and BB) were filtered out by count sum<30. Significance of the result was determined by t value: p<0.1 when t(5,0.95)>2.015 and p<0.05 when t(5,0.975)> 2.571. Ranking differential expressed genes by the significance of the change created lists of genes for further analyses:

*GSEA analysis*

GSEA algorithm was used as described in [6] to evaluate enrichment of CRLF2-IL7RAins gene signatures in Philadelphia and Philadelphia-like cases compared to non-Philadelphia-like cases. Gene expression data for B-ALL patients was obtained from the patient database St. Jude's group (GSE26281). This database included 29 Philadelphia and Philadelphia-like B-ALL cases [BCR-ABL (n=18), CRLF2+ (n=11)] and 98 non-Philadelphia-like B-ALL cases [E2A-PBX (n=8), TEL-AML1 (n=24), MLL rearrangements (n=15), non CRLF2+ Hyperdipliody (n=29), other (n=22)]. Ranked list was generated using free GEO website tool GEO2R.

*Single cell RNA sequencing analysis*

Counts matrix was generated from raw reads using cellranger v.2.1.0[7]. Data was analyzed using Scater[8] package for R as follows: Low quality cells were filtered out by discarding cells that failed either one of these criteria: 1. More than 10% of the cells' detected genes were mitochondrial. 2. Cells had less than 2 median absolute deviations (MADs) of detected genes (less than 455 genes). 3. Cells had less than 2 MADs log10 total counts (less than 728 counts). Next, genes that were expressed in less than 10 cells were also discarded from the counts matrix. T-SNE plots were generated from the filtered counts matrix using the plotTSNE function on a subset of 99 highly variable genes with perplexity set to 20. Differential expression analysis was carried out using edgeR[9] package for R. Genes that obtained absolute fold change greater than 2 and FDR smaller than 0.1 were considered as differentially expressed.

*Whole genome sequencing*

Leukemic and BB transduced corresponding cord blood cells were collected from transplanted mice. gDNA was purified using standard techniques. Sequencing libraries were prepared using NEBNext ULTRA II library preparation kit and sequenced on HiseqXten (BGI Hong Kong).

*SNP array*

Array analysis was done using Affymetrix CytoScan HD array (Affymetrix, California, USA) according to the manufacturer's recommendations (Affymetrix manual protocol Affymetrix® Cytogenetics Copy Number Assay P / N 703038 Rev. 3). The raw data was processed using Chromosome Analysis Suite (ChAS) 3.1.0.15.

1. Hertzberg L, Vendramini E, Ganmore I, et al. Down syndrome acute lymphoblastic leukemia, a highly heterogeneous disease in which aberrant expression of CRLF2 is associated with mutated JAK2: a report from the International BFM Study Group. *Blood*. 2010;115(5):1006-1017.
2. Shochat C, Tal N, Bandapalli OR, et al. Gain-of-function mutations in interleukin-7 receptor-alpha (IL7R) in childhood acute lymphoblastic leukemias. *The Journal of experimental medicine*. 2011;208(5):901-908.
3. Sather BD, Ryu BY, Stirling BV, et al. Development of B-lineage predominant lentiviral vectors for use in genetic therapies for B cell disorders. *Mol Ther*. 2011;19(3):515-525.
4. Tiscornia G, Singer O, Verma IM. Production and purification of lentiviral vectors. *Nat Protoc*. 2006;1(1):241-245.
5. Good Z, Sarno J, Jager A, et al. Single-cell developmental classification of B cell precursor acute lymphoblastic leukemia at diagnosis reveals predictors of relapse. *Nat Med*. 2018;24(4):474-483.
6. Subramanian A, Tamayo P, Mootha VK, et al. Gene set enrichment analysis: a knowledge-based approach for interpreting genome-wide expression profiles. *Proc Natl Acad Sci U S A*. 2005;102(43):15545-15550.
7. Zheng GX, Terry JM, Belgrader P, et al. Massively parallel digital transcriptional profiling of single cells. *Nat Commun*. 2017;8:14049.
8. McCarthy DJ, Campbell KR, Lun AT, Wills QF. Scater: pre-processing, quality control, normalization and visualization of single-cell RNA-seq data in R. *Bioinformatics*. 2017;33(8):1179-1186.
9. Robinson MD, McCarthy DJ, Smyth GK. edgeR: a Bioconductor package for differential expression analysis of digital gene expression data. *Bioinformatics*. 2010;26(1):139-140.

### Supplementary tables

| Gene | Fc | Gene | Fc | Gene | Fc | Gene | Fc | Gene | Fc | Gene | Fc |
|---|---|---|---|---|---|---|---|---|---|---|---|
| IGHG3 | 7.256783893 | IL2RA | 3.578827613 | PIM1 | 2.268236108 | AC005780.1 | 2.832667091 | C10orf10 | 2.755863242 | CARNS1 | 1.693268971 |
| CD36 | 4.50251621 | IGLV6-57 | 3.517932141 | IGLJ2 | 1.255969874 | KLHL17 | 2.245780907 | DNAJB9 | 1.85049231 | HDAC7 | 1.145003348 |
| CRLF2 | 7.122145797 | AC005307.1 | 3.640588628 | CD248 | 2.972288611 | CMTM8 | 2.410090037 | CAV1 | 3.381907207 | DYRK1B | 1.657034962 |
| IGHG1 | 5.750111252 | TMCC2 | 4.11498824 | HIST1H2BJ | 3.332901228 | GNG8 | 1.765510588 | HESX1 | 2.260810393 | SCAMP2 | 1.030481434 |
| IL7R | 6.123840754 | IGHA2 | 5.465483833 | PAQR4 | 2.482666971 | GMPPB | 1.620287611 | TSC22D1 | 1.182773304 | SCCPDH | 1.379524741 |
| SMIM3 | 7.251507754 | LGALS3BP | 2.432153531 | SCN3A | 2.008947143 | CD27 | 2.017019125 | KREMEN2 | 2.005345304 | RP11-452L6.5 | 1.687234551 |
| CCL17 | 6.450762548 | IL15 | 2.635077813 | STX8P1 | 2.26331195 | IGHV4-31 | 2.708913369 | IFI44L | 2.839344159 | STT3A | 1.025061752 |
| PTP4A3 | 2.998927037 | ELFN1 | 3.400094529 | SSR3 | 1.369322759 | MRPL36 | 0.683728874 | ANKRD36BP2 | 1.710172606 | DNAJC3 | 1.360764106 |
| JCHAIN | 2.771991179 | IGKV1D-8 | 3.484656114 | NYNRIN | 2.779862399 | HRSP12 | 1.097716713 | HERPUD1 | 1.333169463 | MEGF6 | 1.651389534 |
| IGHE | 3.356408183 | ENPP2 | 2.633302732 | PPCDC | 1.082079534 | RABAC1 | 1.151486064 | TMEM81 | 2.321072673 | IGHV1-3 | 2.312622459 |
| MAL | 3.772570409 | C19orf54 | 1.768669319 | GOT1 | 1.905799265 | CD70 | 2.183051144 | RP11-588K22.2 | 2.414556647 | HIF1A-AS2 | 1.447576682 |
| GAS6 | 3.68150641 | IGLV10-54 | 2.656158805 | N4BP2 | 1.602322051 | ZMYND11 | 1.065684711 | CD59 | 1.782270171 | IGLV5-48 | 2.172396307 |
| CISH | 4.512570475 | IGLV3-19 | 2.452012993 | KIFC1 | 3.34209993 | AC005519.4 | 1.962502907 | IGLV3-12 | 2.426332512 | ATXN7L2 | 2.27339009 |
| MSC | 3.633770386 | MYDGF | 2.222235205 | MDFIC | 1.375884404 | KLHL9 | 1.586037958 | TMEM187 | 2.10727399 | MYO1B | 1.490504656 |
| IGLV8-61 | 3.244881771 | SOWAHD | 2.996387093 | SDF2L1 | 1.580179983 | SERPINF1 | 1.358826892 | VDR | 1.441365585 | PDXK | 1.024517551 |
| SELM | 3.983929797 | PPP2R3B | 2.24302268 | PRDX4 | 1.704118697 | SOCS1 | 2.287896778 | AC233755.2 | 1.474600773 | SLC35F2 | 0.969041628 |
| IGHG4 | 5.208285447 | HRH1 | 3.858956831 | PIM2 | 1.792667997 | AMBRA1 | 1.70136942 | IGHV3-20 | 1.589806975 | NANS | 0.888717741 |
| SMKR1 | 4.179292693 | HLX | 3.249882862 | LLNLR-268E12.1 | 2.696545739 | MANF | 1.637330565 | SOCS2-AS1 | 2.085272016 | AMN | 1.930033228 |
| CH25H | 4.4028574 | PYCR1 | 2.573710084 | SLC7A5 | 1.648849557 | CCND2 | 1.460788949 | H2AFJ | 1.397729045 | CETN2 | 1.788930816 |
| IGLC7 | 3.746146821 | PLXNB2 | 1.481466959 | IGHV1-2 | 2.206868993 | SLC29A2 | 2.200568618 | P4HB | 1.459340299 | LRRC23 | 2.332741658 |
| LINC00996 | 2.46878752 | CDKN2B | 2.918079971 | ENAM | 2.258587736 | AC007253.1 | 2.507358039 | IGHJ4 | 1.443770297 | E4F1 | 1.435142821 |
| IGHG2 | 6.385957442 | NDFIP2 | 2.84939314 | FAM92B | 2.970266553 | IGHV3-11 | 2.064200217 | GSTP1 | 1.156051896 | KDELR2 | 1.131484213 |
| IGHV3-64 | 3.957132981 | IGLV5-37 | 2.646077049 | TP73 | 2.474875599 | RP11-326C3.11 | 2.203610077 | CYCS | 0.729808641 | MRPL40 | 0.964043602 |
| IGLV4-69 | 1.950712411 | CCR2 | 2.750918874 | ERLEC1 | 1.022108511 | OXER1 | 1.922485035 | SEC61A1 | 1.278287437 | C10orf54 | 1.281573208 |
| TCL1B | 2.654623836 | NINJ1 | 3.002982454 | CTD-2353F22.1 | 2.303976346 | DOK4 | 2.197538016 | AMBN | 2.4348415 | SOCS3 | 3.169679382 |
| IGHA1 | 4.437127892 | IGLV1-47 | 2.332889036 | CASP1 | 1.423863049 | PEBP1 | 1.128453014 | SPCS1 | 0.811174379 | SSR4 | 1.382473731 |
| MAP1A | 4.065960422 | LINC00998 | 0.765123561 | PHLDA3 | 2.498440765 | SDC1 | 3.054940317 | DUSP1 | 1.775282334 | RP11-147L13.14 | 2.2107208 |
| RASD1 | 3.47086839 | TRIB1 | 2.208052221 | BST2 | 1.211731562 | XBP1 | 1.953563032 | HSPA1B | 1.265757912 | RWDD2A | 1.161756162 |
| RGS16 | 3.5138642 | PSAT1 | 2.597481334 | TXNDC11 | 1.592982126 | IGLL1 | 1.483052818 | TLR2 | 2.71246758 | CH17-262H11.1 | 1.70467848 |
| SPAG4 | 2.364928575 | THEMIS2 | 1.922149467 | GADD45G | 2.921329041 | IGLV1-51 | 1.873489012 | SPCS2 | 1.040683206 | SHROOM2P1 | 2.466571847 |
| IGLVI-70 | 3.358129411 | RAG2 | 3.245701312 | AC026202.3 | 2.898918517 | SUSD3 | 1.261267999 | RPSAP36 | 1.55602904 | RP5-855D21.3 | 1.340879141 |
| IGLV4-60 | 2.902266928 | PDIA4 | 1.373780681 | IGHV6-1 | 3.097340786 | NOC4L | 2.747250927 | ARID3B | 0.765890831 | FAM19A5 | 2.34087924 |
| IGLC6 | 2.346819966 | TXNDC5 | 2.078561522 | SNTA1 | 1.97198329 | TOX | 2.664445952 | IFI27L2 | 0.98617297 | GAB1 | 1.290797303 |
| SOCS2 | 4.498353194 | MAN1A1 | 2.159890421 | IGHJ3P | 1.209613405 | CHCHD10 | 1.398906436 | MZB1 | 1.474807733 | TNFRSF17 | 1.145735491 |
| IGKV3D-11 | 2.054119777 | SLC25A23 | 1.826662543 | NOD2 | 1.982329551 | IGHV3-73 | 1.900570492 | SLC35C1 | 1.48550527 | MIR22HG | 1.641200038 |
| NPTX1 | 4.070452685 | TRAT1 | 3.087696181 | RP11-275F13.3 | 2.58690195 | SSH3 | 2.67057052 | ISG15 | 0.881401584 | JAK2 | 1.303898672 |
| RP11-686D22.10 | 2.846083583 | NT5DC2 | 3.0714206 | SELK | 1.437188797 | IGLV7-46 | 1.557576775 | EGR1 | 1.637528749 | SLC38A5 | 1.307527762 |
| PRR5L | 2.942285422 | IFNAR2 | 1.21383764 | H1F0 | 2.727178266 | GZMB | 1.631253411 | MLLT4 | 1.517845867 | PDF | 1.84654494 |
| SLFN12L | 2.364514043 | CCDC136 | 2.833124889 | C10orf128 | 1.832164951 | SAV1 | 1.004642298 | IGHV2-26 | 2.310604003 | TAS2R64P | 1.776506104 |
| TMEM121 | 3.291948594 | WFS1 | 2.805522454 | POLR3D | 0.958410747 | RP1-45N11.1 | 1.488916229 | TIGD6 | 1.653552504 | SLAIN1 | 1.413099676 |
| RP11-1094M14.5 | 2.989354932 | RAPGEF3 | 3.434865571 | HYOU1 | 1.367324562 | TNFRSF4 | 2.49604589 | FAM120AOS | 0.905850555 | HAPLN3 | 2.536967629 |
| IGLV7-43 | 4.025949593 | RP11-524N5.1 | 1.716788093 | BCL2L1 | 1.663996371 | FAS | 1.85389269 | PPIB | 1.163605352 | CDR2 | 1.321582553 |
| CA6 | 3.460596071 | FAM46C | 1.710003498 | IGLV1-36 | 2.792856983 | ARF4 | 0.692076057 | AURKAIP1 | 0.800858014 | GNAQ | 2.28258407 |
| GUCY1A3 | 2.880756863 | RAG1 | 3.400821779 | HSPA1A | 2.059418371 | DLEU7-AS1 | 2.814869811 | GAS7 | 1.38262963 | NECAB3 | 2.454783985 |
| IGHGP | 4.478777646 | TEX22 | 1.999384015 | HSPA5 | 1.426261324 | MME | 1.673674086 | TMED2 | 1.150402138 | CHST15 | 1.509264624 |
| BHLHA15 | 3.1556109 | IGLC3 | 2.267726725 | PREB | 1.307167057 | FKBP2 | 1.045711628 | CALR | 1.249074076 | LL22NC03-88E1.18 | 1.800715492 |
| IFITM1 | 2.375616228 | DGCR6L | 1.191559813 | ITM2C | 1.079152657 | IFITM2 | 0.840136268 | RCBTB2 | 1.865159868 | KSR1 | 0.843124825 |
| IGHV3-30 | 4.212092113 | DANCR | 1.174079592 | RP11-1070N10.3 | 2.6975662 | CDKN2A | 1.380188165 | CAPN2 | 0.942727663 | MRPL24 | 0.930883535 |
| CHPF | 3.681222676 | LST1 | 1.207164504 | PLSCR1 | 1.365952922 | DHRS9 | 1.358892194 | PRAMENP | 2.121196262 | DNAJB5 | 2.478513439 |
| MGAT3 | 4.051940085 | NET1 | 1.24245622 | LARGE-AS1 | 2.551665589 | PDCD1LG2 | 2.346735142 | TMEM217 | 2.406731158 | IGF2 | 1.915207071 |

**Supplementary table 1:** Top differentially over expressed genes in CRLF2-IL7RAins vs backbone transduced engrafted cells. Genes are ranked according to the significance of the change (t). Fc – average of mean $Log_2$Fold change of paired CRLF2-IL7RAins over Backbone transduced engrafted cells from same cord blood batch.

| IL-6 JAK-STAT3 signaling | mTORC1 signalling | B-lymphocyte progenitir | myc max targets | appoptosis reversed by IL6 | apoptosis | p53 pathway | unfplded protein response |
|---|---|---|---|---|---|---|---|
| CD36 | PSAT1 | IL7R | CISH | PTP4A3 | CASP1 | RGS16 | PSAT1 |
| CRLF2 | GOT1 | TCL1B | CH25H | GAS6 | H1F0 | PLXNB2 | WFS1 |
| IL2RA | SDF2L1 | SOCS2 | RGS16 | RGS16 | BCL2L1 | CDKN2B | SLC7A5 |
| PIM1 | SLC7A5 | RAG2 | PYCR1 | PIM1 | CCND2 | NINJ1 | HYOU1 |
| SOCS1 | HSPA5 | WFS1 | TRIB1 | PIM2 | FAS | CASP1 | HSPA5 |
| FAS | XBP1 | RAG1 | HIST1H2BJ | HSPA1A | CAV1 | PHLDA3 | PREB |
| TLR2 | FKBP2 | SCN3A | SDF2L1 | PLSCR1 | DNAJC3 | SOCS1 | XBP1 |
| SOCS3 | CALR | GADD45G | HRSP12 | SOCS1 | IGF2R | CCND2 | DNAJB9 |
| CXCL13 | SERP1 | CD27 | MANF | C10orf10 | CD69 | SDC1 | HERPUD1 |
| TNFRSF1B | GAPDH | IGLL1 | SAV1 | DNAJB9 | CDKN1A | FAS | SPCS1 |
| BAK1 | HSP90B1 | MME | FKBP2 | SOCS3 | BCL2L11 | CDKN2A | CALR |
| IL10RB | CXCR4 | C10orf10 | TMEM187 | CHST15 | GCH1 | TSC22D1 | DNAJC3 |
| CD38 | RPN1 | CAV1 | CALR | MSMO1 | FDXR | VDR | SERP1 |
| IL7 | CDKN1A | MZB1 | MRPL40 | CXCR4 | BMF | H2AFJ | HSP90B1 |
| STAT1 | GLRX | CHST15 | SERP1 | HSPA13 | GADD45A | SEC61A1 | PDIA6 |
| JUN | BHLHE40 | IGF2R | HSP90B1 | MX1 | BCL2L2 | CDKN1A | SPCS3 |
| CSF1 | CANX | CITED2 | ST6GALNAC4 | KIAA0125 | ISG20 | CCND3 | CEBPB |
| | EPRS | HIST1H4H | FKBP11 | BCL2L11 | MGMT | SLC3A2 | YIF1A |
| | PDK1 | CD69 | HIST1H2BE | BHLHE40 | CD38 | FDXR | MTHFD2 |
| | FDXR | CXCR4 | RRAS | GADD45A | TAP1 | PERP | SSR1 |
| | ENO1 | GLRX | DCTN4 | CEBPB | BCAP31 | GADD45A | EIF2AK3 |
| | MTHFD2 | CACNB3 | TMED10 | IRF4 | DAP | BAK1 | IFIT1 |
| | SSR1 | HIST1H3H | SLC39A8 | POU2AF1 | MCL1 | TRIAP1 | |
| | ALDOA | P4HA2 | PDK1 | MT1X | CASP3 | PPP1R15A | |
| | PPP1R15A | REXO2 | SLC39A7 | STAT1 | IFITM3 | TAP1 | |
| | LDHA | IRF4 | RBM15B | MCL1 | JUN | ZNF365 | |
| | PGK1 | WSB2 | YIF1A | OAS1 | PSEN1 | BAIAP2 | |
| | UNG | AEBP1 | SNRPA | PRDM1 | PPP2R5B | JUN | |
| | BCAT1 | ISG20 | H2BFS | JUN | | EPHA2 | |
| | PSMD14 | UXS1 | KPNA3 | TMEM184B | | HINT1 | |
| | PSMD13 | POU2AF1 | LDHA | | | PMM1 | |
| | PRDX1 | SMAD1 | EIF3I | | | CD81 | |
| | YKT6 | LMF1 | MAT2B | | | | |
| | USO1 | EIF2AK3 | VCP | | | | |
| | TPI1 | CTBP2 | SPTLC2 | | | | |
| | HSPE1 | PKD2 | CUTA | | | | |
| | WARS | VSIG10 | TESK2 | | | | |
| | CACYBP | JUN | PSMD14 | | | | |
| | FADS2 | CSGALNACT1 | AP4M1 | | | | |
| | IDI1 | TCF3 | HIST2H2BE | | | | |
| | | VPREB1 | SAP30 | | | | |
| | | CD19 | DMAP1 | | | | |
| | | ELK3 | SEC61B | | | | |
| | | LRIG1 | TIMM22 | | | | |
| | | VDAC1 | ARL1 | | | | |
| | | ING1 | WDR4 | | | | |
| | | SEMA6A | PSMD8 | | | | |
| | | | USO1 | | | | |
| | | | HSPE1 | | | | |
| | | | POLE3 | | | | |
| | | | MYBBP1A | | | | |
| | | | SARS | | | | |
| | | | CDK16 | | | | |
| | | | CACYBP | | | | |
| | | | GMDS | | | | |
| | | | DDB1 | | | | |
| | | | PTGES3 | | | | |
| | | | PSMB7 | | | | |
| | | | AZIN1 | | | | |
| | | | NDUFA1 | | | | |
| | | | ING1 | | | | |

**Supplementary table 2:** Top leading genes in GSEA. Ranked list of differentially expressed genes from CRLF2-IL7RAins vs Backbone engrafted cells were analyzed for gene set enrichment (GSEA algorithm, Broad institute). Top genes leading to enrichment plot are listed.

| Genomic Position | Tumor Call | Effect |
|---|---|---|
| chr1:6128316-6550966 | ./.:1:0 | DEL |
| chr2:88855243-88914861 | ./.:0:0 | DEL |
| chr2:88916530-89021436 | ./.:1:0 | DEL |
| chr2:89022486-89032415 | ./.:0:0 | DEL |
| chr2:89034492-89055526 | ./.:1:0 | DEL |
| chr2:89056678-89082665 | ./.:0:0 | DEL |
| chr2:89085229-89088777 | ./.:1:0 | DEL |
| chr2:89089906-89108033 | ./.:2:0 | DEL |
| chr2:89109152-89244680 | ./.:1:0 | DEL |
| chr2:89246139-89250404 | ./.:2:0 | DEL |
| chr2:89251664-89298147 | ./.:1:0 | DEL |
| chr2:89940432-90095478 | ./.:1:0 | DEL |
| chr3:138798511-138808252 | ./.:1:0 | DEL |
| chr3:82584435-82600604 | ./.:3:1 | DUP |
| chr7:38256528-38329654 | ./.:0:0 | DEL |
| chr7:38331081-38361383 | ./.:1:0 | DEL |
| chr9:137532898-137572364 | ./.:1:0 | DEL |
| chr9:5633386-37485713 | ./.:1:0 | DEL |
| chr14:105866868-106032920 | ./.:3:0 | DUP |
| chr14:106034543-106153593 | ./.:2:0 | DEL |
| chr14:106155125-106405332 | ./.:3:1 | DUP |
| chr14:106514372-106554068 | ./.:3:1 | DUP |

Within 88825371-90316061 IGK locus

Within 105536746-106879844 IGH locus

**Supplementary table 3:** Leukemia genomic structural variations. Leu*kemic* cells and batch matched backbone transduced engrafted cord blood (representing germline) were sequenced (whole genome sequencing 60x). The table depicts major structural variations between the samples.

| type | gene | chr | pos | Risk allele (frequency) | dbsnp | Genotype* |
|---|---|---|---|---|---|---|
| intronic | GATA3 | 10 | 8062245 | A (0.19) | rs3824662 | 0/1:23,22:45 |
| intronic | ARID5B | 10 | 61963818 | C (0.35) | rs10821936 | 0/1:18,12:30 |
| intronic | ARID5B | 10 | 61992400 | G (0.37) | rs7089424 | 0/1:21,18:39 |
| UTR5 | CEBPE | 14 | 23119522 | C (0.66) | rs2239635 | 0/1:26,18:44 |
| ncRNA_exonic | CDKN2B-AS1 | 9 | 22029548 | C (0.18) | rs564398 | 0/1:25,15:40 |

**Supplementary table 4:** list of SNPs in leukemic sample: List of SNPs that were found in both leukemic sample and CB-matched backbone transplanted sample; chr: chromosome, pos: position, Risk allele (frequency): SNP in sample and frequency in general population according to 1000Gemones, Genotype explanation: 0/1 = het 11,20= depth of each allelle,:31 overall depth.

| pre-leukemia vs BB | Leukemia vs BB | CRLF2-IL7RAins vs BB |
|---|---|---|
| LST1 | SOCS2 | CRLF2 |
| XBP1 | SPATS2L | GAS6 |
| SOCS2 | XBP1 | CISH |
| SPATS2L | LST1 | SOCS2 |
| CD99 | CD99 | NPTX1 |
| KCNA5 | IFITM2 | CA6 |
| MME | KCNA5 | IFITM1 |
| SH3BP5 | ECM1 | IL2RA |
| CCND2 | TUBA4A | ENPP2 |
| NCF2 | S100A4 | RAPGEF3 |
| IFITM2 | MME | LST1 |
| CR2 | NCF2 | MDFIC |
| DCTN4 | NPTX1 | CASP1 |
| CD34 | GIMAP4 | CD27 |
| TNFRSF1B | CISH | AMBRA1 |
| ENG | | CCND2 |
| DUSP26 | | DOK4 |
| IFITM3 | | XBP1 |
| LPAR6 | | MME |
| S100A4 | | IFITM2 |
| CISH | | CD59 |
| IFITM1 | | MYO1B |
| ADGRE5 | | PDXK |
| GAS6 | | CD69 |
| ECM1 | | SLC2A5 |
| EFNA1 | | ST6GALNAC4 |
| SEMA6A | | GBP1 |
| IL2RA | | TNFRSF1B |
| ADGRG1 | | CALHM2 |
| ANGPT2 | | DCTN4 |
| RBM47 | | ITGA6 |
| DENND3 | | SLC48A1 |
| GIMAP4 | | GADD45A |
| FYN | | CLEC2B |
| | | ENG |
| | | ADGRE5 |
| | | ROR1 |
| | | ZNF711 |

**Supplementary table 5:** Top leading genes in Ph-like signature by GSEA**.** A gene-list of Ph-like signature was generated based on publicly available data base (St. Jude's group (GSE26281)), using the free GEO website tool- GEO2R. The list was used for gene set enrichment analysis (GSEA- broad institute) of pre-leukemia, leukemia and CRLF2-IL7Rains, compared with backbone. Top leading genes are listed.

| Antigen | Fluorochrome | Manufacturer |
|---|---|---|
| CD45 | Vio Green | Miltenyi |
| | APC | Biolegend |
| CD123(IL7RA) | APC-Alexa 700 | Beckman Coulter |
| | BV421 | Biolegend |
| CD19 | ECD | Beckman Coulter |
| | APC-Alexa750 | Beckman Coulter |
| IgM | APC | eBioscience |
| CD10 | PC7 | Beckman Coulter |
| CD34 | APC | BD |
| CD16/32 | none | Biolegend |
| Nucleic acid | 7AAD | BD |
| LIVE/DEAD fixable Dead cells staining | Near IR/Violet | ThermoFisher Scientific (Molecular probes) |

**Supplementary table 6** Antibodies and markers for flow cytometry.

| Protein | Clone | Manufacturer | Metal Isotope | Staining |
|---|---|---|---|---|
| 4EBP1(pT36/T46) | 236B4 | Cell Signaling Technology | Nd144 | Intracellular |
| Akt (pS473) | D9E | Cell Signaling Technology | Tb159 | Intracellular |
| BTK (pY551/511) | 24A/BTK | BD Biosciences | Yb174 | Intracellular |
| cCaspase3 | C92-605 | BD Biosciences | Ho165 | Intracellular |
| CD10 | HI10a | Biolegend | Gd156 | Surface |
| CD127 | A019D5 | Biolegend | Dy162 | Surface |
| CD16 | 3G8 | Fluidigm | Bi209 | Surface |
| CD179a | HSL96 | Biolegend | Sm149 | Intracellular |
| CD179b | HSL11 | Biolegend | Gd158 | Intracellular |
| CD19 | H1B19 | Biolegend | Nd142 | Surface |
| CD20 | 2H7 | Biolegend | Sm147 | Surface |
| CD22 | HIB22 | Biolegend | Nd143 | Surface |
| CD235 | HIR2 | Biolegend | In115 | Surface |
| CD24 | ML5 | Biolegend | Gd160 | Surface |
| CD3 | UCHT1 | Biolegend | Er170 | Surface |
| CD34 | 581 | Biolegend | Nd148 | Surface |
| CD38 | HIT2 | Biolegend | Er168 | Surface |
| CD43 | CD43-10G7 | Biolegend | Er167 | Surface |
| CD45 human | HI30 | Fluidigm | Y89 | Surface |
| CD45 mouse | 30F11 | Biolegend | In113 | Surface |
| CD79b | CB3-1 | Biolegend | Nd146 | Surface |
| cPARP | F21-852 | BD Biosciences | La139 | Intracellular |
| Creb (pS133) | 87G3 | Cell Signaling Technology | Yb176 | Intracellular |
| CRLF2 | 1A6 | eBioscience | Dy161 | Surface |
| CyclinA (total) | BF-683 | BD Biosciences | Sm154 | Intracellular |
| CyclinB1 (total) | GNS-1 | BD Biosciences | Dy164 | Intracellular |
| Erk1/2 (pT202/pY204) | D13-14-4E | Cell Signaling Technology | Yb173 | Intracellular |
| Glucocorticoid Receptor | D8H2 | Cell Signaling Technology | Eu151 | Intracellular |
| GFP | SF12.4 | Fluidigm | Tm169 | Intracellular |
| HistoneH3 (pS28) | HTA28 | Biolegend | Ce140 | Intracellular |
| IgHintracellular | polyclonal | Novus | Eu153 | Intracellular |
| IgH surface | MHM-98 | Fluidigm | Yb172 | Surface |
| Ikaros (total) | D10E5 | Cell Signaling Technology | Nd145 | Intracellular |
| Ki67 | B56 | BD Biosciences | Sm152 | Intracellular |
| PU.1 | 9G7 | Cell Signaling Technology | Gd157 | Intracellular |
| RB (pS807/811) | J112-906 | BD Biosciences | Er166 | Intracellular |
| rpS6 (pS235/pS236) | N7-548 | BD Biosciences | Lu175 | Intracellular |
| SRC (pY418) | K98-37 | BD Biosciences | Pr141 | Intracellular |
| STAT5 (pY694) | 47 | BD Biosciences | Gd155 | Intracellular |
| Syk (pY319/pY352) | 17a | BD Biosciences | Yb171 | Intracellular |
| TdT | E17-1519 | BD Biosciences | Dy163 | Intracellular |

**Supplementary table 7:** Antibodies and reagents for mass cytometry.

#### Supplementary figures

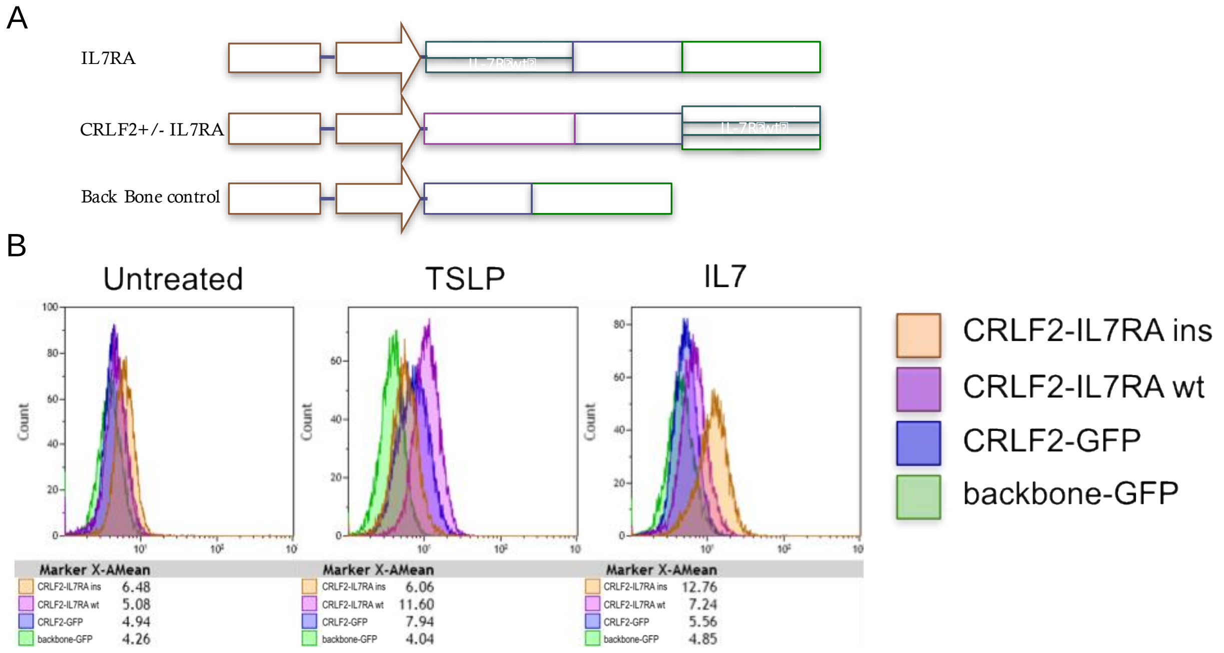

**Supplementary figure 1** *CRLF2 and IL7RA over expression* (A) Diagram of pRRL Eµ B29 B-cell vector used for expression of GFP and bi-cistronic expression of combinations of CRLF2, IL7RAwt, IL7RAins ppcl. with GFP. (B) Flow cytometer histogram of phospho STAT5 in transduced 018Z (BCP-ALL) cells after activation with TSLP (2ng/ml) or IL7 (2ng/ml). Mean fluorescent intensities are portrayed in the histogram and values are listed below.

A

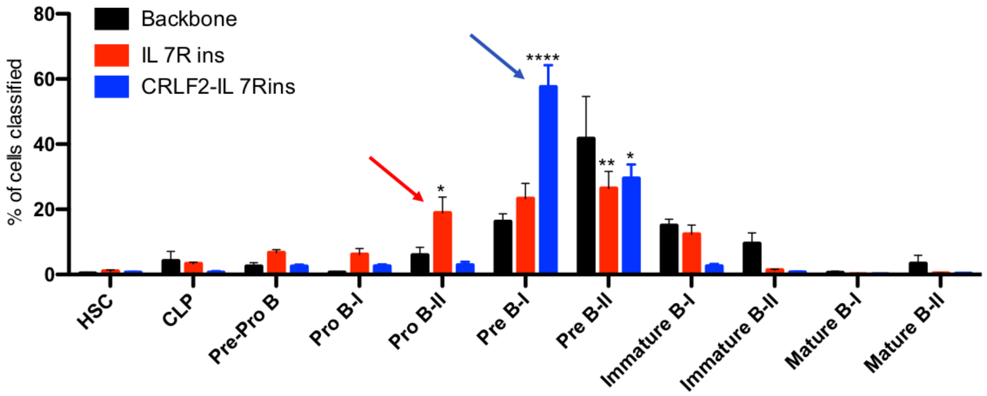

B

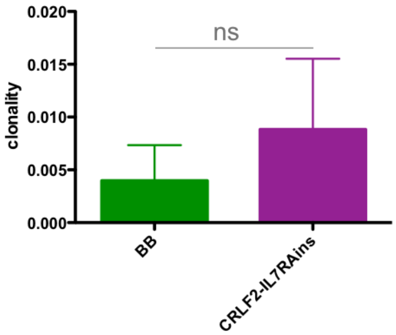

**Supplementary figure 2**: Early B-cell differentiation and clonality in CRLF2/IL7RAins and control backbone transduced cells. A) Human cells from BM of engrafted mice were analyzed by mass cytometer. Bar-graph represents mean percentage of cells from: backbone (n=3), IL7Rins (n=4) and CRLF2-IL7Rins (n=3). Samples classified in each B-lineage developmental stage by using the developmental classifier. Statistical analysis was done by two-way ANOVA followed by Tukey test for multiple comparison corrections. The asterisks indicate statistically significant difference compare to the backbone group (*p<0.05, **p<0.01, ****p<0.0001). Arrows indicate early B-cell precursor accumulation. B) Bar-graph representing total sample clonality of BM CD10$^+$ and CD19$^+$ sorted cells from CB batch paired BB and CRLF2 IL7RAins transplanted mice. Bars are mean +/- SEM of n=3 paired transplanted transduced CB. Statistical analyses were performed using paired two tailed t-test.

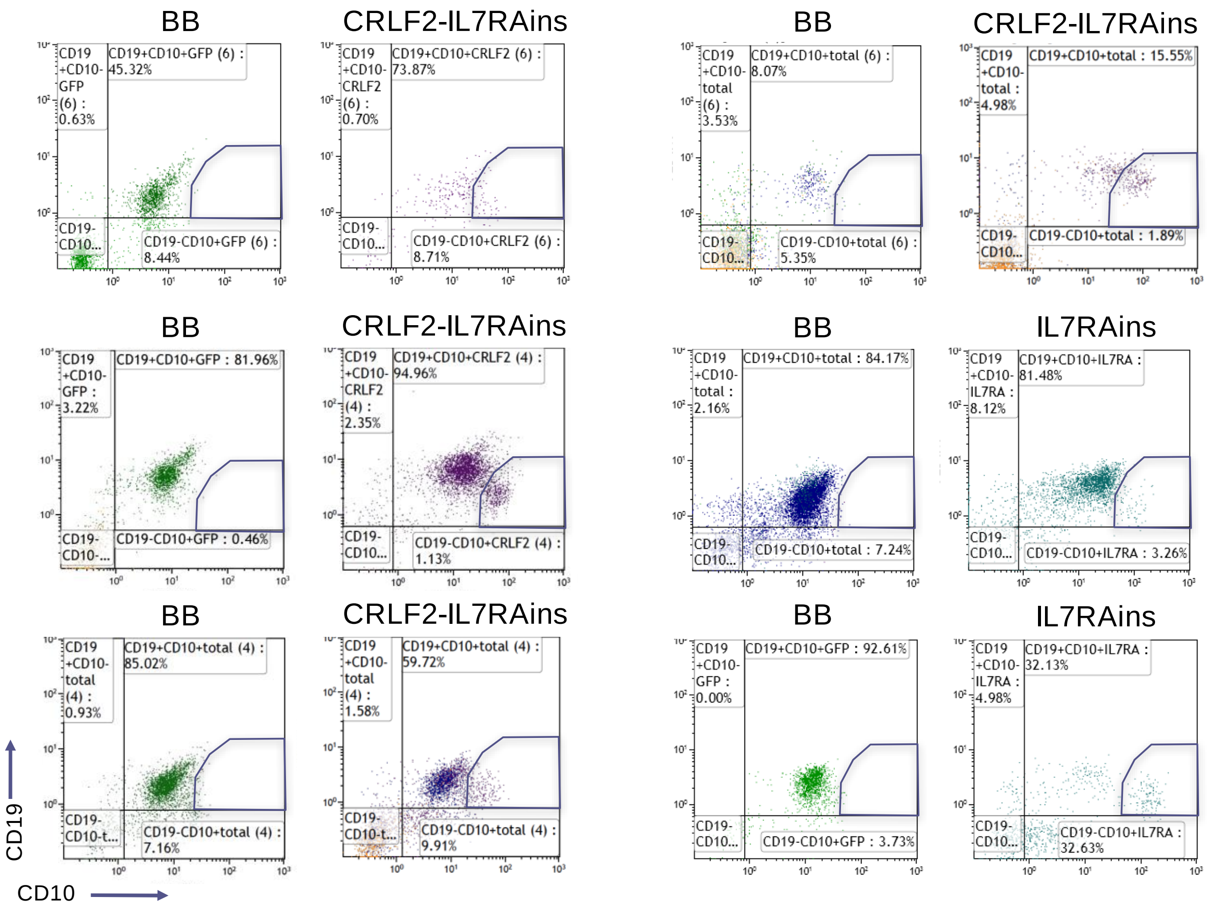

**Supplementary figure 3:** Unique CD10highCD19low population in IL7RAins transduced cells. Flow cytometry immunophenotyping of engrafted backbone and CRLF2-IL7RAins or IL7RAins transduced cells. Paired samples are CB batch matched.

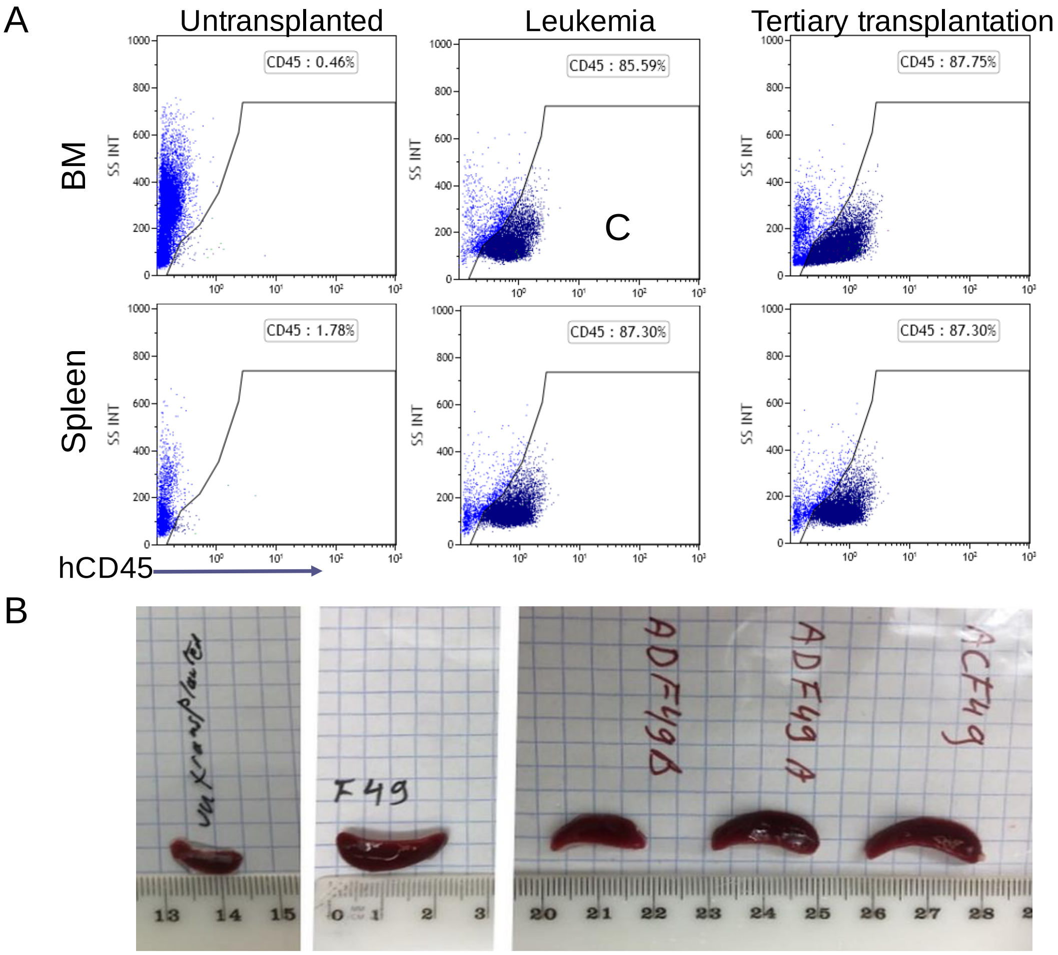

**Supplementary figure 4:** Human engraftment in leukemic mice. A) Flow cytometry charts of BM and spleen of untransplanted mouse (left) secondary IL7RA engrafted mouse that developed leukemia (center) and tertiary transplanted mouse that was engrafted with cells from spleen of leukemic mouse (right) B) Pictures depicting spleen size at sacrifice of untransplanted mouse (left) secondary IL7RA engrafted mouse that developed leukemia (center) and three tertiary transplanted mouse that were engrafted with cells from spleen of leukemic mouse (right).

V-(D)-J rearrangement summary for query sequence (multiple equivalent top matches, if present, are separated by a comma):

| Top V gene match | Top D gene match | Top J gene match | Chain type | stop codon | V-J frame | Productive | Strand |
|---|---|---|---|---|---|---|---|
| IGHV3-15*01,IGHV3-15*02 | IGHD3-10*01 | IGHJ4*02 | VH | Yes | Out-of-frame | No | + |

V-(D)-J junction details based on top germline gene matches:

| V region end | V-D junction* | D region | D-J junction* | J region start |
|---|---|---|---|---|
| ACAGA | TGGGGCGC | TACTATGGTTCGGGGAGTTATTATAAC | TCCAT | ACTAC |

*: Overlapping nucleotides may exist at V-D-J junction (i.e, nucleotides that could be assigned to either rearranging gene). Such nucleotides are indicated inside a parenthesis (i.e., (TACAT)) but are not included under the V, D or J gene itself.

Sub-region sequence details:

| | Nucleotide sequence | Translation | Start | End |
|---|---|---|---|---|
| CDR3 | ACCACAGATGGGGCGCTACTATGGTTCGGGGAGTTATTATAACTCCATACTACTTTGACTAC | TTDGALLWFGELL*LHTTLTT | 226 | 287 |

Alignments

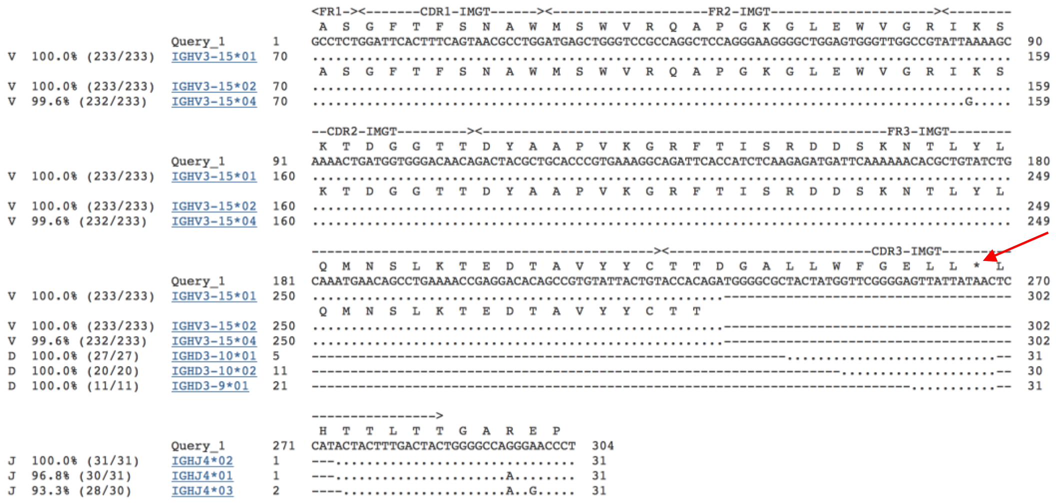

**Supplementary figure 5:** Non-functional rearrangement of leukemia cells. NCBI- BLAST query output of the leukemia IGH sequence. Arrow pointing to the stop codon.

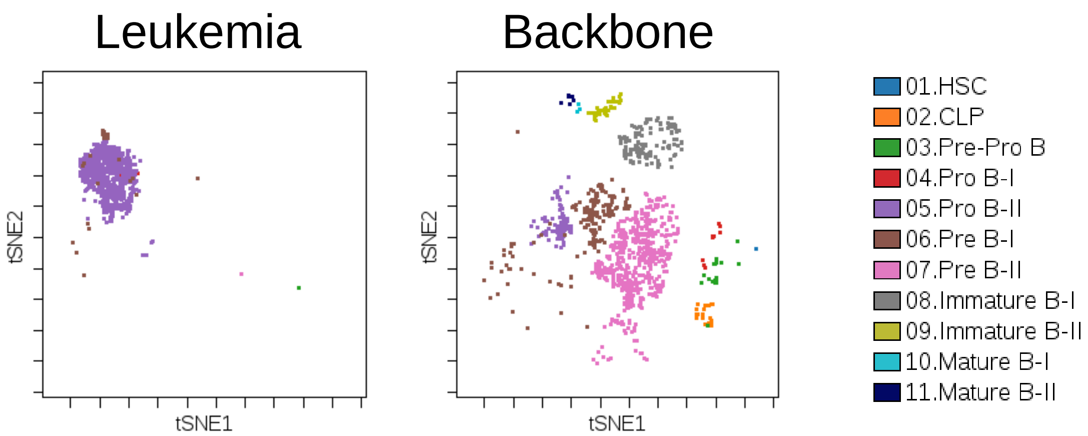

**Supplementary figure 6:** B-cell differentiation analysis by mass cytometry. tSNE maps of leukemic cells and engrafted cells from backbone transduced matched CB. B-cell developmental stage was analyzed by mass cytometer using developmental classifier.

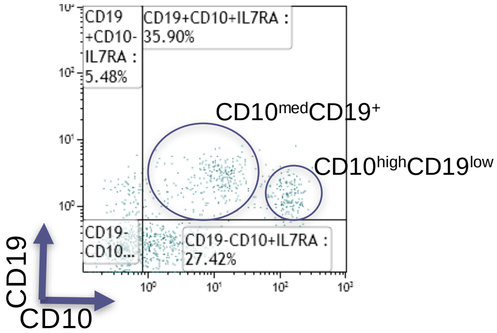

| Sample | Est Total Nucleated Cells | Fraction Nucleated | nucleotide | copy | Copy Normalized | count | frequency | Frequency Normalized |
|---|---|---|---|---|---|---|---|---|
| CD10_med CD19+_50K | 19241.73 | 0.000185638 | TGAACAGCCTGAAAACCGAGGACACAGCCGTGTATTACTGTACCACAGATGGGGCGCTACTATGGTTCGGGGAGTTATTATAACTCCATACTACTTTGACTACTGGGGCCAGGGAACC | 113 | 19 | 3 | 0.020047404 | 0.020757995 |
| CD10highCD19low_4539 | 957.89 | 0.000734322 | TGATTCAAAAAACACGCTGTATCTGCAAATGAACAGCCTGAAAACCGAGGACACAGCCGTGTATTACTGTACCACAGATGGGGCGCTACTATGGTTCGGGGAGTTATTATAACTCCATACTACTTTGACTACTGGGGCCAGGGAACC | 150 | 4 | 1 | 0.079933495 | 0.094854162 |

**Supplementary figure 7:** Frequency of pre-leukemic clone in sorted populations from primary mouse. Viably frozen cells from the primary mouse from which the leukemic clone was developed were thawed and analyzed. Top: flow cytometry plot with marked sorted populations. Bottom table summarizing the reported frequencies of leukemic rearrangement after VH-region sequencing of genomic DNA from the sorted populations.

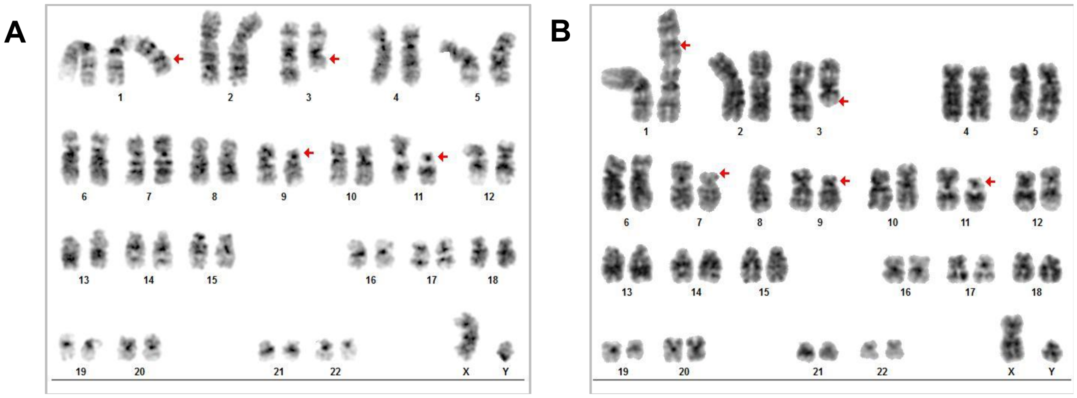

**Supplementary figure 8:** Karyotype analysis of leukemic cells**.** Leukemic cells (15 metaphases) from spleen and BM of tertiary transplanted mice were analyzed by G-banding karyotype analysis. Major clone (A) and a sub-clone (B) karyotypes with significant chromosomal aberrations (as pointed by red arrows) are:
46,XY,add(1)(p32),del(3)(q24),del(9)(p13),del(11)(p11.2)[9]/46,idem,del(7)(p13)[6].
.

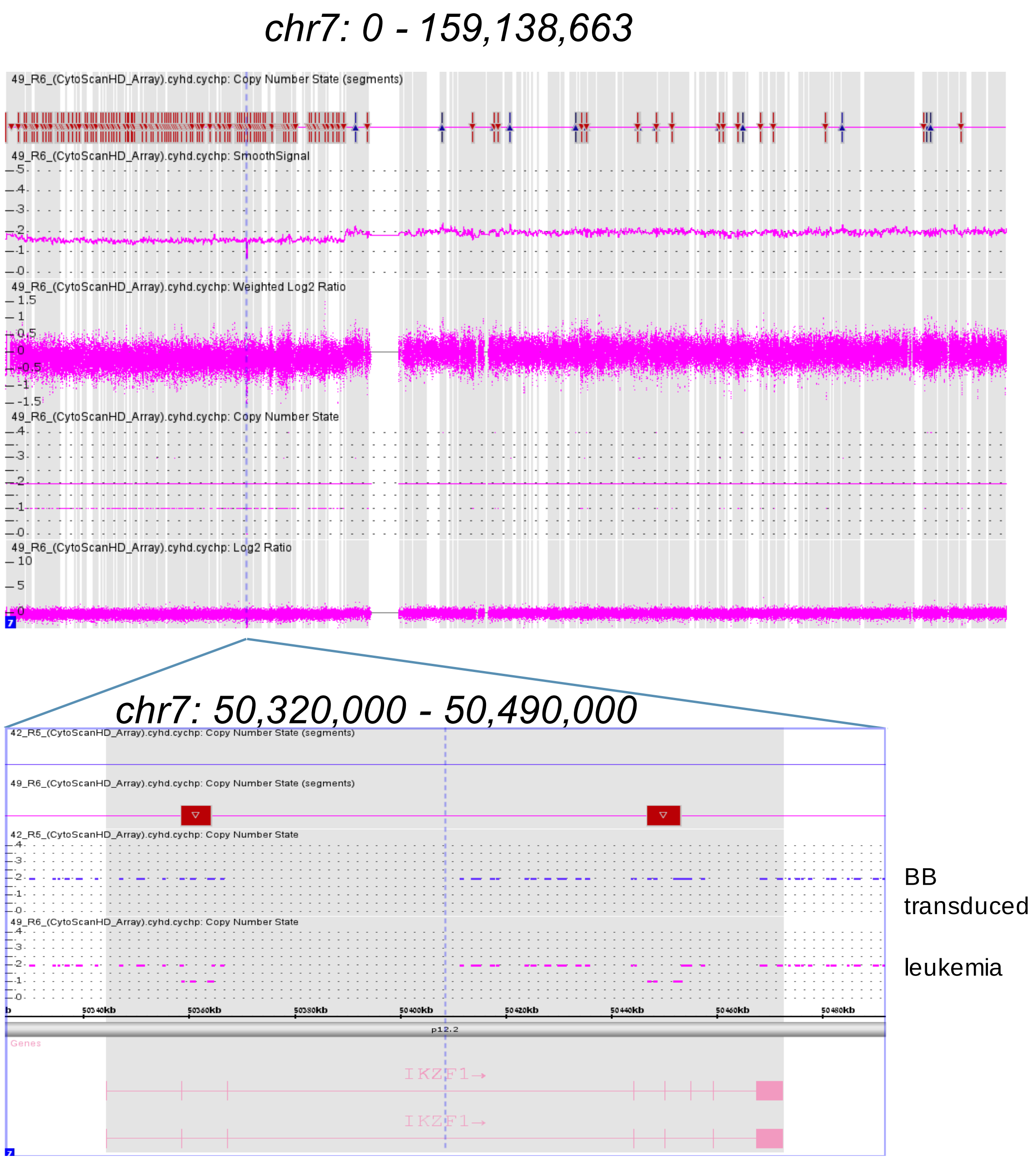

**Supplementary figure 9:** IKZF1 internal deletions in leukemic cells: Leukemic cells and corresponding backbone transduced and transplanted cord blood were subjected to SNP array analysis. Image depicts deletions in chromosome 7 encompassing IKZF1 region. Upper: SNP array analysis depicting chromosome 7 in whole revealing deletions around/surrounding p12.1 – p22.3. Lower: SNP array analysis focusing on IKZF1 Genomic region depicting pronounced deletion around exons 2,5 in leukemia but not in Germline (BB-transduced cord blood).

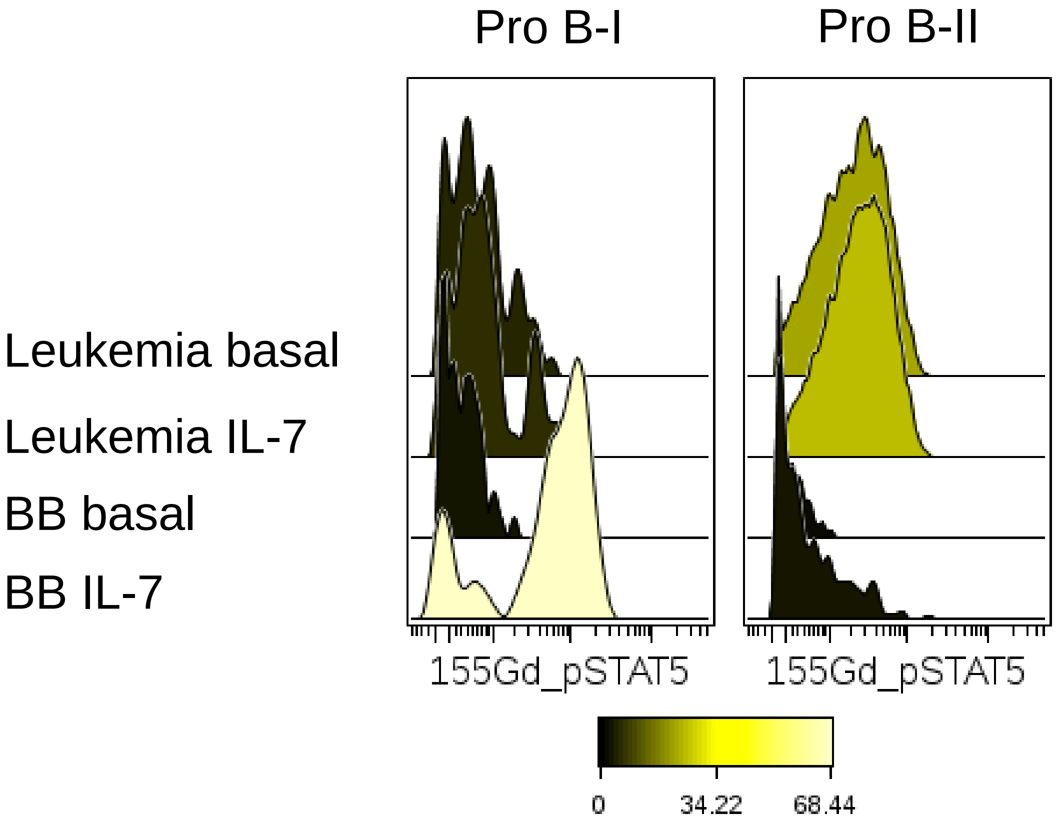

**Supplementary figure 10:** Cytokine independent activation of JAK-STAT signaling in leukemic cells. Histograms representing mass cytometer analysis of pSTAT5 with and without IL7 activation (100ng/ml) of ProB-I and ProB-II Leukemic cells and engrafted BB transduced cells from matched CB.

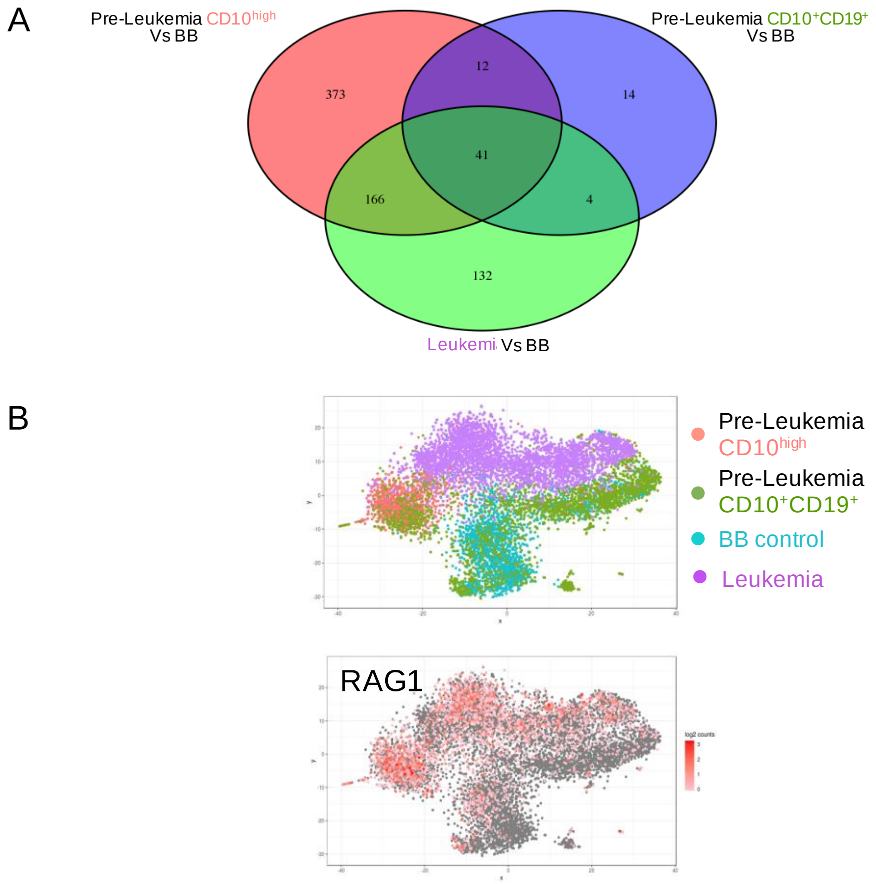

**Supplementary figure 11:** scRNAseq of leukemic and pre-leukemic populations. (A) Venn diagram of differentially expressed genes in bulk analysis of pre-leukemia and leukemia samples vs BB control sample (B) Top: Transcriptome correlation t-SNE map after 10X scRNAseq, bottom: Relative expression of selected genes displayed on t-SNE map
